# Thylakoid grana stacking revealed by multiplex genome editing of LHCII encoding genes

**DOI:** 10.1101/2021.12.31.474624

**Authors:** Zeno Guardini, Rodrigo L. Gomez, Roberto Caferri, Johannes Stuttmann, Luca Dall’Osto, Roberto Bassi

## Abstract

Land plant chloroplasts differ from algal ones for their thylakoid membranes being organized in grana: piles of vesicles paired by their stromal surface, forming domains including Photosystem (PS) II and its antenna while excluding PS I and ATPase to stroma membranes, connecting grana stacks. The molecular basis of grana stacking remain unclear. We obtained genotypes lacking the trimeric antenna complex (Lhcb1-2-3), the monomeric Lhcb4-5-6, or both. Full deletion caused loss of grana, while either monomers or trimers support 50% stacking. The expression of Lhcb5 alone restored stacking at 50%, while Lhcb2 alone produced huge grana which broke down upon light exposure. Cyclic electron transport was maintained in the lack of stacking, while excitation energy balance between photosystems and the repair efficiency of damaged Photosystem II were affected. We conclude that grana evolved for need of regulating energy balance between photosystems under terrestrial canopy involving rapid changes in photon spectral distribution.

## Introduction

The land plant cell chloroplast contains chlorophylls (Chl) and carotenoids (Car) within its inner membrane system called thylakoids, as recognized already by Meyer in 1883 (Gunning et al., 2006). Its most distinctive feature is the thylakoid organization into grana stacks (Wietrzynski et al., 2020), a feature of land plant species, with size dependent on growth light intensity (Heitz, 1936) and spectra (Iwai et al., 2018; Kyle et al., 1983), which is absent in algae (Vallon et al., 1991). Grana are stacks of round, flattened vesicles (Granick & Porter, 1947) connected to each other by pairs of unstacked membranes, called stromal membranes, intersecting grana at an angle, suggesting a helical arrangement (Paolillo & Reighard, 2011). Freeze-etching electron microscopy (EM) coupled to mutant analysis showed that the photosynthetic pigment-protein complexes of thylakoid membranes are compartimented: photosystem (PS) II particles in the appressed region of the grana (Armond & Arntzen, 1977; Simpson & Robinson, 1984); PSI and ATPase complexes hosted in stromal membranes (Armond et al., 1977; Simpson, 1982). Cytochrome (Cyt) *b*_6_*f* complex was enriched in stromal domains and grana margins (Allred & Staehelin, 1985; Olive et al., 1986; Vallon et al., 1991). Biochemical and immunocytochemical analysis supported differential compartimentation of PSI and PSII (Anderson & Melis, 1983; Vallon et al., 1991), arguably for preventing excitation energy spillover from PSII to PSI, owing to the low energy chlorophyll spectral forms of plant PSI-LHCI (Murata, 1969). Since photosynthetic electron transport occurs linearly between PSs, the Plastoquinone (PQ) diffusion towards Cyt *b*_6_*f* is restricted by PSII crowding in grana partitions (Lavergne & Joliot, 1991) similar to the diffusion of plastocyanin (PC) in the narrow thylakoid lumen (Kirchhoff et al., 2011), thus limiting linear electron flow (LET) rate. Restriction of diffusion also applies to PSII damaged by photoinhibition, which need to reach the stromal domains in order to be repaired (Mattoo et al., 1989). Thus, the size of granal *vs*. stromal domains appear to compromise between the positive effect of ensuring adequate excitation energy to both PSs and the negative effect of restricting PQ diffusion and PSII repair rate. This is consistent with PQ overreduction activating STN-7 and -8 kinases, which phosphorylates, respectively, the Lhcb2 subunit trimeric antenna (LHCII) and PSII core, yielding into reduced granal diameter (Bassi et al., 1989; Hepworth et al., 2021). Thus, LHCII subunits, diffuse to PSI-LHCI. This boosts PSI photon cross section causing oxidation of plastoquinol (PQH_2_) (Kyle et al., 1983) and kinase inactivation, balancing PSI *vs*. PSII activity. It was proposed that the reduction of grana domains enhances the fraction of Cyt *b*_6_*f* in stromal membranes and favors cyclic electron flow (CEF) and ATP synthesis, thus helping to meet the NADPH/ATP ratio required for CO_2_ fixation (Kramer & Evans, 2011) and adding to the regulatory value of grana stacking dynamics. In this context it is important to clarify the mechanism of grana stacking. Reverse genetics and EM analysis yielded controversial results: intermittent light grown plants becomes LHCII-deficient and exhibit limited grana (Argyroudi-Akoyunoglou et al., 1971; Armond et al., 1976); however, *chlorina f2* mutants of barley lack Chl *b* and are depleted in LHCs, yet they retained extensive grana stacks and photosystem lateral heterogeneity (Bassi et al., 1985; Kim et al., 2009). Grana stacks were also retained in the *viridis115* barley mutants, lacking PSII core complex, and in the double mutant *viridis 115 x chlorina f2* (Simpson et al., 1989), suggesting that neither of the two major thylakoid components located in grana partitions (LHCII and PSII core) was essential for grana stacking. Chloroplasts lacking grana can be found in bundle sheath (BS) cells of maize leaves, with a reduced PSII activity and a low LHCII level (Bassi et al., 1995). Also, LHCII of BS exhibited a simpler polypeptide composition with respect to mesophyll chloroplasts (Bassi & Simpson, 1986) suggesting that specific gene products within LHC, rather than any member of the family, might be responsible for stacking. Here, we proceeded to the reverse genetic analysis of grana stacking by a combination of genome editing (GE) and EM. Previous work showed that PSII core complex-less plants plants (Belgio et al., 2014; Campoli et al., 2009; Margulies, 1966) enhance stacking, we focused on LHCs: genotypes lacking either monomeric or trimeric LHC gene products or both were produced and analyzed for stacking by transmission EM. In addition, Lhcb2, a minor component trimeric LHCII, reported to be needed for state 1 - state 2 transitions (Pietrzykowska et al., 2014), was expressed as the only LHC component and found to induce the formation of extensive stacking in dark conditions, which were broken down upon light-induced phosphorylation. Our results show that a major thylakoid stacking is caused by minority components of the PSII antenna system, namely Lhcb5 (CP26) and Lhcb2, which respectively mediate the constitutive and dynamic components of thylakoid stacking, which is required for efficient of PSII repair and balance of excitation energy; instead, it does not significantly change the rate of linear *vs*. cyclic electron transport.

## Results

### Construction of *Arabidopsis* genotypes with selectively reduced PSII antenna system by GE

The construction of *Arabidopsis thaliana* genotypes affected in LHC protein composition was performed from either the knock-out (ko) line *koLhcb3*, missing the Lhcb3 component of the major LHCII complex, or the *NoM koLhcb3* genotype, devoid of the *Lhcb4-6* genes encoding monomeric LHCs: Lhcb4 (CP29), CP26 and Lhcb6 (CP24)(Dall’Osto et al., 2017), as well as Lhcb3 (Supplementary Figure S1-A). To obtain the Lhcb-free genotype, CRISPR-CAS9 targeted mutagenesis strategy was implemented according to (Ordon et al., 2020): for each of the 5 LHCB1 and 3 LHCB2 genes, pairs of sg (single guide) RNAs were designed within coding sequences (Supplementary Table S1), with the aim of producing large DNA deletions, favoring DNA repair by non-homologous end-joining (NHEJ) (Malzahn et al., 2017). *Lhcb1* and *Lhcb2* genes were separately targeted to get Lhcb1-less and Lhcb2-less plants, which were then crossed to obtain lines lacking both Lhcb1 and Lhcb2 proteins. Depending on whether this procedure was applied on *koLhcb3* or *NoM koLhcb3* genotype, we obtained either the *koLHCII* line (lacking all LHCII but retaining monomeric Lhcb4-6 proteins), or the *koLhcb* line, lacking both the monomeric and trimeric LHCs of PSII. In both cases LHCI proteins were unaffected. The *low-LHCII* genotype was selected for the level of residual LHCII among the products of incomplete gene editing of both *Lhcb1* and *Lhcb2*: the two selected lines contain one LHCII trimer per monomeric PSII core complex. *NoM koLhcb1 koLhcb3* lines (*Lhcb2-only*) retained Lhcb2 as the unique PSII antenna (Supplementary Figure S1-B, Supplementary Table S2).

### Growth and pigment composition of LHC-depleted plants

Plants of the different genotypes are shown in Figure 1A upon growth under light-limiting (LL) conditions (150 µmol photons m^-2^ s^-1^, 8/16h photoperiod, 24 °C). Wild type showed the best growth. *NoM* and the two *koLHCII* lines accumulated between 45% and 50% with respect to the wild type; while the growth of *koLhcb* was far more reduced: the two independent lines grew less than 3% with respect to the wild type based on fresh weight (FW) (Figure 1B). The Chls content per leaf surface displayed a behavior similar to the growth pattern although changes were far smaller: *NoM* showed 25% less Chl *vs*. the wild type, while *koLhcb* lines about 50% Chls when compared to *koLHCII* and 25% with respect to the wild type, suggesting that the growth rate per pigment unit was higher in the presence of monomeric antenna proteins (Figure 1C). The pigment composition of the different genotypes reflected the abundance of the pigment-protein complexes residual from gene deletions. Thus, Chl *a*/*b* ratio was the lowest in *NoM* (3.28 vs 3.51 in the wild type), the highest in *koLhcb* (6.21) and intermediate in *koLHCII* (5.19), consistent with the lower Chl *b* content of monomeric Lhcbs vs LHCII (Table I). LHCI complexes, unaffected in our genotypes, account for the residual Chl *b* content in *koLhcb*. The Chl/Car ratio ranged from 3.0 to 3.7 slightly decreasing with LHC content (Table I). The higher Car /Chl ratio in *koLHC* lines resulted from increased β-carotene accumulation, which doubled in *koLhcb* with respect to the LHCII-rich wild type and *NoM* (Supplementary Table S3). Neoxanthin decreased by 60% and 80%, respectively in *koLHCII* and *koLhcb*, with a concomitant increase in violaxanthin content owing to the preferential location of neoxanthin in trimeric LHCII and its absence in PSI-LHCI complexes (Schiphorst & Bassi, 2020).

**Figure 1.**
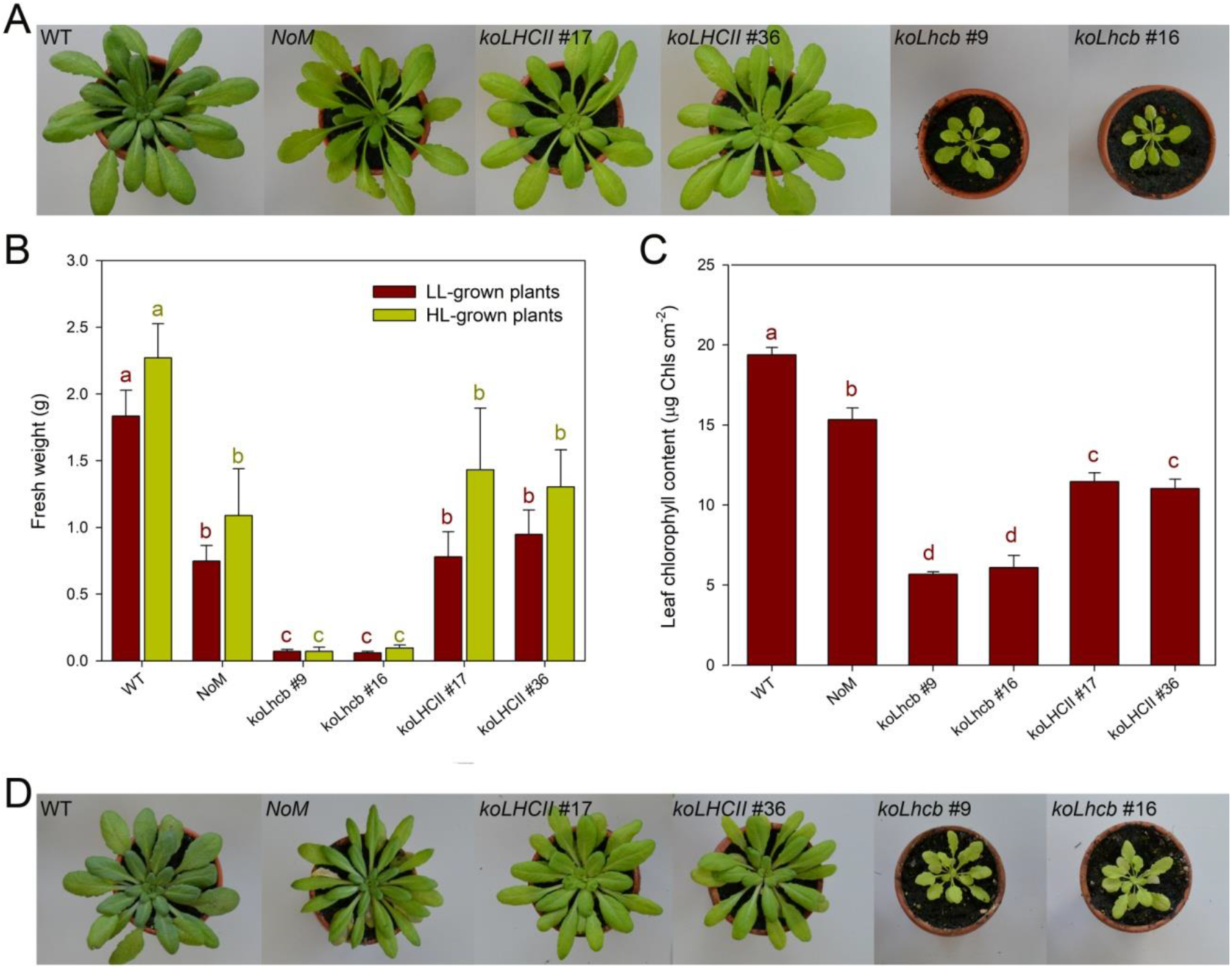
Phenotype of wild type and mutant plants. **(A)** Plants were grown for 6 weeks at 150 µmol photons m^−2^ s^−1^, 23 °C, 8/16 h light/dark (LL). **(B)** The fresh weight of all. Values are expressed as mean ± SD, n=10. **(C)** Leaf Chl content. *koLhcb* lines retained only 25% of Chl per area, while *NoM* and *koLHCII* retained 75% and 50%, respectively. Values are expressed as mean ± SD, n=4. **(D)** Plant growth for 6 weeks at 350 µmol photons m^−2^ s^−1^, 23 °C, 8/16 h light/dark (HL). *NoM* and *koLHCII* showed similar growth while *koLhcb* lines were much smaller, alike in CTRL light (panel B). Leaf Chl content relative to the wild type remained essentially the same in LL vs HL (panel C). Values that are significantly different from the corresponding wild type (ANOVA followed by Tukey’s post-hoc test at a significance level of P < 0.05) are marked with different letters.

**Table 1.** Pigment content and fluorescence induction parameters determined for leaves of *Arabidopsis* WT and mutant lines. At least five different plants were tested for each line. Chl/Car, molar ratio between chlorophylls (*a* + *b*) and carotenoids. Fresh weight refers to growth after 6 weeks under control conditions. Data are expressed as mean ± SD, *n* = 6 biologically independent leaves. Values marked with different letters are significantly different from each other within the column (ANOVA, followed by Tukey’s post-hoc test at a significance level of *P* < 0.05). Experiments were repeated independently twice, with similar results.

|  | Chl a / b | Chl / Car | µg Chl cm<sup>-2</sup> | F<sub>v</sub>/F<sub>m</sub> |
| --- | --- | --- | --- | --- |
| **WT** | 3.51 ± 0.09<sup>a</sup> | 3.69 ± 0.21<sup>a</sup> | 19.4 ± 0.5<sup>a</sup> | 0.82 ± 0.01<sup>a</sup> |
| **NoM** | 3.28 ± 0.05<sup>a</sup> | 3.45 ± 0.09<sup>a,b</sup> | 15.3 ± 0.74<sup>b</sup> | 0.61 ± 0.01<sup>b</sup> |
| **koLHCII #17** | 5.23 ± 0.11<sup>b</sup> | 3.37 ± 0.06<sup>b</sup> | 11.46 ± 0.55<sup>c</sup> | 0.76 ± 0.01<sup>c</sup> |
| **koLHCII #36** | 5.15 ± 0.10<sup>b</sup> | 3.34 ± 0.02<sup>b</sup> | 11.03 ± 0.58<sup>c</sup> | 0.76 ± 0.01<sup>c</sup> |
| **koLhcb #9** | 6.28 ± 0.15<sup>c</sup> | 3.12 ± 0.05<sup>c</sup> | 5.68 ± 0.15<sup>d</sup> | 0.50 ± 0.02<sup>e</sup> |
| **koLhcb #16** | 6.14 ± 0.22<sup>c</sup> | 3.09 ± 0.08<sup>c</sup> | 6.10 ± 0.76<sup>d</sup> | 0.54 ± 0.03<sup>e</sup> |
| **ch1** | - | 2.77 ± 0.02<sup>c</sup> | 6.55 ± 0.54<sup>d</sup> | 0.78 ± 0.01<sup>c</sup> |
| **ch1 koLhcb5** | - | 2.57 ± 0.02<sup>c</sup> | 4.50 ± 0.23<sup>e</sup> | 0.67 ± 0.01<sup>d</sup> |
| **lowLHCII #14** | 4.01 ± 0.22<sup>d</sup> | 3.01 ± 0.06<sup>b</sup> | 7.24 ± 0.87<sup>c</sup> | 0.52 ± 0.05<sup>e</sup> |
| **lowLHCII #32** | 4.71 ± 0.39<sup>b,d</sup> | 2.98 ± 0.03<sup>b</sup> | 7.58 ± 1.01<sup>c</sup> | 0.55 ± 0.05<sup>e</sup> |
| **Lhcb2-only** | 3.96 ± 0.16<sup>e</sup> | 3.18 ± 0.11<sup>b</sup> | 9.77 ± 1.82<sup>c</sup> | 0.51 ± 0.02<sup>e</sup> |

The major differences in growth between mutants might be due to either the large differences in PSII antenna size, thus limiting the photon harvesting, and/or to defects in the electron transport chain induced by changes in the organization of the thylakoid membranes. We thus proceeded to grow plants at higher irradiance, expected to compensate for antenna size limitation, yet avoiding excess irradiation in order to circumvent photodamage. Figure 1D shows wild type and LHC mutant genotypes grown at moderately high light (HL, 350 µmol photons m^-2^ s^-1^, 8/16h photoperiod, 24°C). *NoM* and *koLHCII* grew similarly while *koLhcb* was far slower and grew as in control light light (150 µmol photons m^-2^ s^-1^) (Figure 1B, D). Thus, enhancing irradiance did not complement for the decreased growth phenotype evidenced under LL conditions and, in some cases, made it even stronger. This suggests that insufficient photon harvesting was unlikely the major cause of the differences in growth between genotypes. On the other hand, none of the lines were obviously photodamaged, or died at 350 µmol photons m^-2^ s^-1^, suggesting that photoprotection was stronghly reduced.

### Pigment-protein complexes and photosynthetic function in antenna mutants

The actual pigment-protein content of the genotypes as well as their supramolecular organization was investigated by non-denaturing Deriphat-PAGE upon solubilization with low (0,8%) α-DM. Consistent with previous reports (Dall’Osto et al., 2014, 2017), the *NoM* genotype was depleted in PSII supercomplexes and over-accumulated the major trimeric antenna LHCII (Figure 2A). The trimeric LHCII band was completely absent in both *koLHCII* and *koLhcb* genotypes. These were also depleted of high molecular weight (MW) supercomplexes except for two bands migrating just above PSI-LHCI (BD1 and BD2). A faint green band (BD3) was detected migrating slightly below trimeric LHCII in both *koLHCII* and *koLhcb*, while a fourth band (BD4) was detected in both *koLHCII* and *koLhcb*, migrating as the monomeric Lhcbs (Figure 2A). Biochemical and spectroscopical analyses revealed BD1 and BD2 being composed of PSI-LHCI complexes with different LHC complements, BD3 contained dimeric LHCI subunits, BD4 comprised monomeric Lhcbs in *koLHCII* and monomeric LHCI in *koLhcb* (Supplementary Figure S2).

**Figure 2.**
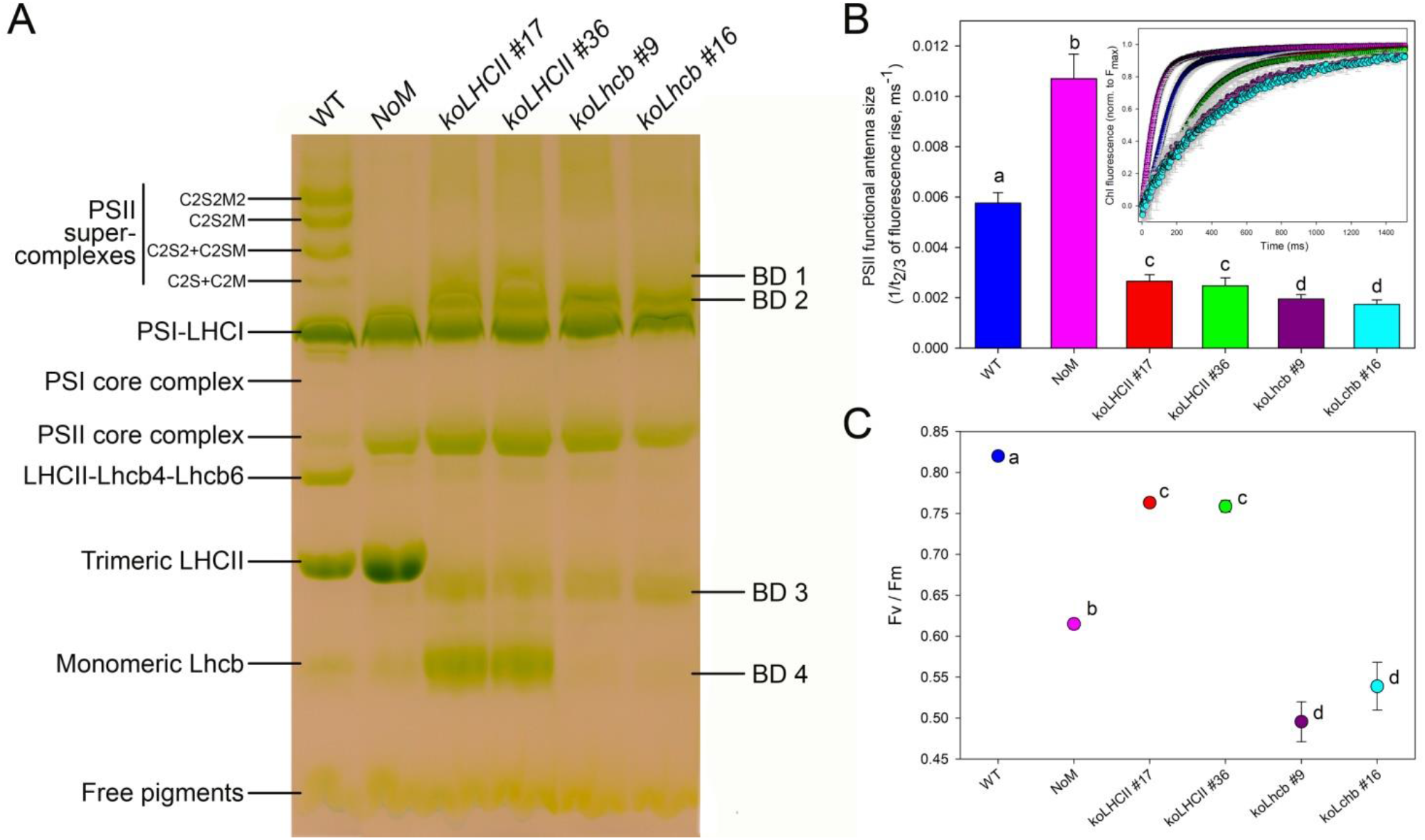
Biochemical and functional characterization of the photosynthetic apparatus of wild type and mutant plants. **(A)** Non-denaturing Deriphat-PAGE of thylakoids upon solubilization with 0.8% α-DM, revealing pigment-protein complexes of wild type, *NoM, koLHCII* and *koLhcb* lines. Thylakoid proteins corresponding 20-35 µg of Chls were loaded in each lane. The composition of major bands is indicated based on previous reports, while that of bands BD1-BD4 was determined from absorption spectra and SDS-PAGE (Supplementary Figure S2). **(B)** PSII functional antenna size of wild type and mutants, measured at RT on leaves vacuum-infiltrated with 50 μM DCMU. The reciprocal of time corresponding to two-thirds of the fluorescence rise (t_2/3_^-1^) was taken as a measure of the PSII functional antenna size. Plants were dark-adapted for 30 min before measurements. Data are expressed as mean ± SD (n=9). Values marked with different letters are significantly different from each other (ANOVA followed by Tukey’s post-hoc test at a significance level of P < 0.05). **(C)** PSII maximal quantum yield (F_v_/F_m_) of wild type, *NoM, koLHCII* and *koLhcb* lines, measured on dark adapted leaves. Symbols and error bars show means ± SD (n=4). Values that are significantly different (ANOVA followed by Tukey’s post-hoc test at a significance level of P < 0.05) are marked with different letters.

The functional antenna size of PSII was determined from the rise time of Chl fluorescence in DCMU-treated leaves. Compared to the wild type, *NoM* had nearly twice the capacity for PSII photon harvesting, while *koLHCII* and *koLhcb* antenna size, scored 30% and 45% respectively, *vs*. the wild type (Figure 2B). When PSII activity was probed without DCMU in order to determine the maximum quantum yield of photochemistry, the wild type and *koLHCII* scored very high, i.e. 0.82 and 0.76 respectively. Instead, *NoM* and *koLhcb* showed a low quantum yield, corresponding to 0.62 and 0.51 respectively (Figure 2C).

### Thylakoid membranes organization

Antenna proteins of PSII are the most abundant components of thylakoid membranes, suggesting that thylakoid organization could be affected in LHC-less genotypes. Indeed, the trimeric LHCII complex has been suggested to be the major determinant of grana stacking (Standfuss et al., 2005) and of the consequent domain segregation of the thylakoids, which is typical of land plant chloroplasts and is the basis for multiple regulation and biogenetic mechanisms essential for photosynthesis in the challenging terrestrial environment, such as PSII repair cycle and state 1 - state 2 transitions. To verify the effect of LHCII abundance in the different genotypes, we have analyzed the extent of grana stacks by transmission EM (Figure 3A-G). All genotypes had chloroplasts with the same size in ultrathin sections (Figure 3H). Grana stacks were defined as made of at least three thylakoid appressions; the stacks diameter was approx. 450 nm and was found to be very similar between genotypes, only *NoM* showed a slightly larger grana diameter of approximately 550 nm (Figure 3I). However, see below for the special case of the *Lhcb2-only* genotype.

**Figure 3.**
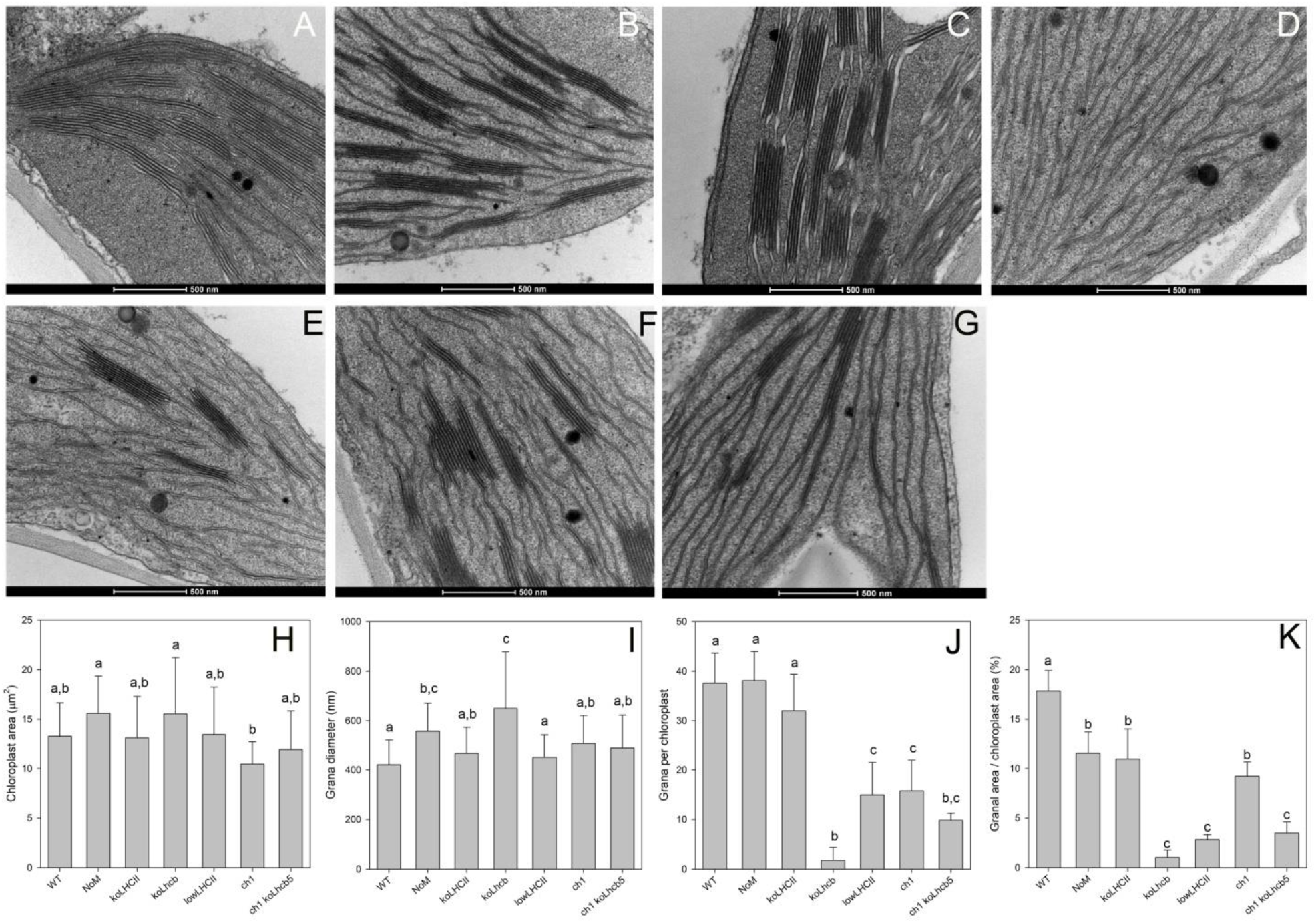
Transmission electron micrographs of plastids from leaf mesophyll cells of wild type and mutant lines. **(A-D)** Plants grown in short-day conditions were dark-adapted for 2 hours before harvesting leaves, then samples were fixed, embedded, and observed in thin sections at different levels of magnification. Wild type (A), *NoM* (B) and *koLHCII* (C) showed a characteristic organization of stroma lamellae interconnecting grana, while the chloroplasts of *koLhcb* (D) lacked grana. **(E-G)** Micrograph of *lowLHCII* (E), *ch1* (F) and *ch1 koLhcb5* (G) chloroplasts. **(H-K)** Statistical analysis of the extent of the thylakoid stacking in the wild type, LHC KO lines and Chl *b*-less mutants *ch1* and *ch1 koLhcb5*. Histograms report: (H) chloroplast area (n≥12); (I) grana diameter [n = 22 (wild type), 27 (*NoM*), 26 (*koLHCII, Lhcb2-only*), 7 (*koLhcb*), 25 (*ch1* lines); (J) grana per chloroplast (n≥8); (K) granal area *vs*. chloroplast area (n≥5). Data are shown as mean ± SD. Values that are significantly different (ANOVA followed by Tukey’s post hoc test at a significance level of P < 0.05) are marked with different letters.

Quantification of grana stack was performed by measuring the surface of grana stack *vs*. chloroplast area, since the total length of thylakoid membranes was th same across genotypes, (Figure 3K). Wild type chloroplasts had about 18% of their section area occupied by grana, a value that was reduced to 11% in *koLHCII* and *NoM*. Deletion of LHCII in *koLhcb* caused a virtual absence of grana stacks (<1% of stacked area, Figure 3K), while partial depletion of LHCII in the *lowLHCII* genotype yielded only a very partial recovery on surface occupied by grana to 3%. We notice that *koLHCII* retains more than 50% of its stacking ability, similar to *NoM* (Figure 3K), even though these two genotypes display highly diverse content in LHCs: *koLHCII* has three monomeric LHCs per PSII core, while *NoM* has about 6 LHCII trimers per PSII core, i.e. 60% more LHCII with respect to the wild type (Dall’Osto et al., 2017). When comparing the stacking efficiency for a single LHC monomeric unit, the score is 6 times higher than when comparing the wild type to *NoM*, 3 times higher than comparing *koLHCII* to *lowLHCII*, the latters having the same LHC/PSII core stoichiometry. This suggests that, although both monomeric and trimeric LHCs contributed to stacking, the monomeric LHCs were most critical determinants for grana formation. In all cases the number of events analyzed was statistically significant except for the case of *koLhcb*, because most of the chloroplasts did not contain any stacks (Figure 3J).

Possibly, one of the clearest effects on grana stacking in previous research was obtained by controlling the expression of CURT genes (Heinz et al., 2016). Supplementary Figure S3 shows that no major differences were observed in the abundance of CURT, suggesting LHCII abundance/composition controlled stacking independently form CURT, or LHCII was needed for CURT activity.

To identify whether a specific subunit, among monomeric LHCs, had a special role in stacking, we analyzed the *ch1* genotype devoid of CAO (Chl *a* oxygenase), which is known for a strongly reduced antenna size (Kim et al., 2009) caused by Lhcb proteins being destabilized in lack of Chl *b*. CP26, however, is particularly promiscuous for Chl *a vs*. Chl *b* binding (Croce et al., 2002) and is retained and functional in *ch1* genotype (Havaux et al., 2007). We have analyzed by EM the extent of grana stacking in *ch1* and *ch1 koLhcb5* mutants (Figure 3F-G). First, we assessed that the chloroplast area and the diameter of grana in sections were very similar in *ch1* and *ch1 koLhcb5* genotypes with respect to the wild type and *koLHCII* (Figure 3H,I). *ch1* had similar stacking as the *koLHCII* genotypes (Figure 3K), suggesting that whatever induced stacking in *koLHCII* was still present in *ch1*. Instead, stacking was drastically reduced in the *ch1 koLhcb5* genotype (Figure 3G,K), implying that CP26, among monomeric LHCs, had a prominent function on grana formation.

### Lhcb2 controls changes in grana stacking during State 1 - State 2 transitions

*Arabidopsis* lines depleted on Lhcb2 proteins lacked state 1 - state 2 transitions (Pietrzykowska et al., 2014). The GE of *Lhcb1* in the *NoM koLhcb3* background (Supplementary Figure S1) produced a genotype retaining Lhcb2 as the only PSII antenna (Ordon et al., 2020). Wild type and mutant genotypes analyzed above showed grana diameters between 400 and 550 nm (Figure 3I), (Kirchhoff, 2019). The *Lhcb2-only* plants were striking different with respect to the *lowLHCII* lines: Despite having a similar PSII-core/LHCII ratio (Supplementary Figure S1), they also showed a few, yet much larger, grana with a diameter up to 3 µm or even more (Figure 4A-C), spanning the chloroplast sections, besides those alike observed wild type. We then proceeded to verify whether light exposure did affect thylakoid stacking. Dark-adapted leaves were illuminated with PSII light (200 µmol photons m^-2^ s^-1^, 24°C, 60 min) in order to promote PQ over-reduction despite the small PSII antenna size of both *Lhcb2-only* and *lowLHCII* genotypes, thus Lhcb2 phosphorylation (Figure 4D). The very in *Lhcb2-only* chloroplasts large grana essentially broke down upon illumination, and the size distribution extended towards narrower diameters. In both *Lhcb2-only* and *lowLHCII* lines, grana length and level of stacking were significantly reduced by state 2 induction (Figure 4E-F).

**Figure 4.**
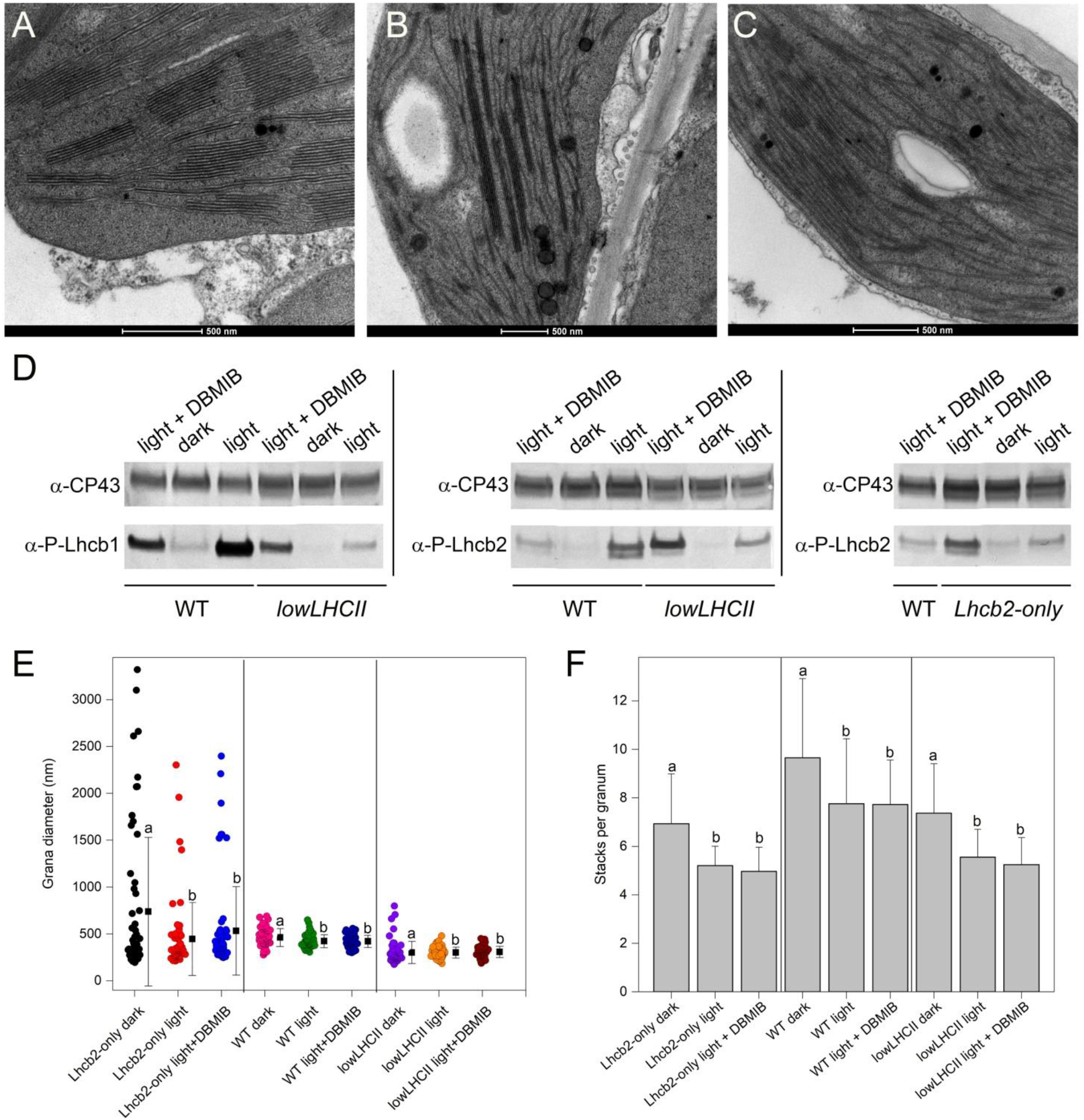
Changes in chloroplast ultrastructure of state 1- and state 2-adapted leaves. **(A-C)** Leaves from wild type (A), *Lhcb2-only* (B) and *lowLHCII* (C) plants were subjected to 2 hours of PSII light (200 µmol photons m^−2^ s^−1^) to promote LHCII phosphorylation and transition to state 2. Reference samples included leaves either dark-adapted for 1 hours or exposed to PSII light upon vacuum-infiltration with 100 μM DBMIB to maximize PQ reduction thus LHCII phosphorylation. Leaf discs were then fixed, embedded, and observed in thin sections. **(D)** Leaf discs from the same treatments were snap frozen, then the phosphorylation level of Lhcb1/Lhcb2 was quantified by immunotitration, using α-lhcb1-P and α-lhcb2-P primary antibodies. Proteins corresponding to 0.8 µg of Chls (wild type sample) or 2.4 µg of Chls (mutant samples) were loaded for each sample. All samples were loaded on the same SDS-PAGE slab gel. **(E)** Statistical analysis of differences in grana diameter, in state 1 and state 2 [n = 60 (*Lhcb2-only*), 55 (wild type), 58 (*lowLHCII*)]. **(F)** Statistical analysis of the average number of layers in grana stacks, in state 1 and state 2 (n = 30). Data (E, F) are expressed as mean ± SD. Values that are significantly different (ANOVA followed by Tukey’s post-hoc test at a significance level of P < 0.05) are marked with different letters.

### Physiological consequences of thylakoid stacking for photosynthesis

Chloroplasts of unicellular algae do not show well-defined grana stacks, while cyanobacteria exhibit single membranes, showing that oxygenic photosynthesis can proceed without the need of a stacked organization of thylakoids (Mullineaux, 2005). Nevertheless, consequences of decreased level of stacking are clear from Figures 1 and 3: growth is dependent on the level of stacking, with the wild type showing both the maximal level of membrane stacking and the best growth. *NoM* and *koLHCII* exhibited intermediate level of both biomass yield and thylakoid stacking, while *koLhcb* has no grana and was severely impaired in growth. While these data clearly showed that growth rate and grana stacking were related, the functional reason(s) for the growth phenotype are still unclear. Grana stacking has been suggested to affect (1) light-harvesting, (2) the switch between CEF and LEF and/or (3) the repair of damaged PSII (Hepworth et al., 2021; Koochak et al., 2019; Mullineaux & Emlyn-Jones, 2005; Pribil et al., 2014). We proceeded to evaluate whether the PSII repair rate was affected by thylakoid stacking. To this aim, leaves were treated with excess light to induce photoinhibition, as detected by F_v_/F_m_ ratio, which was reduced to about 30% of initial values in all genotypes. The recovery of F_v_/F_m_ was then followed under low light conditions (15 µmol photons m^-2^ s^-1^, 24°C) favoring the PSII repair process (Mattoo et al., 1999). The results are shown in Figure 5A. Clearly, the wild type and *koLHCII* genotypes showed the same rate of PSII recovery. A similar dataset was published previously for the *NoM* genotype (Dall’Osto et al., 2020), showing no difference in the kinetic of recovery with respect to wild type plants. Instead, *koLhcb* healed its PSII activity to a much lower extent, suggesting that lateral heterogeneity between fully functional PSII in grana and PSII under repair in stroma-exposed margins might be necessary for a fully functional PSII repair cycle.

**Figure 5.**
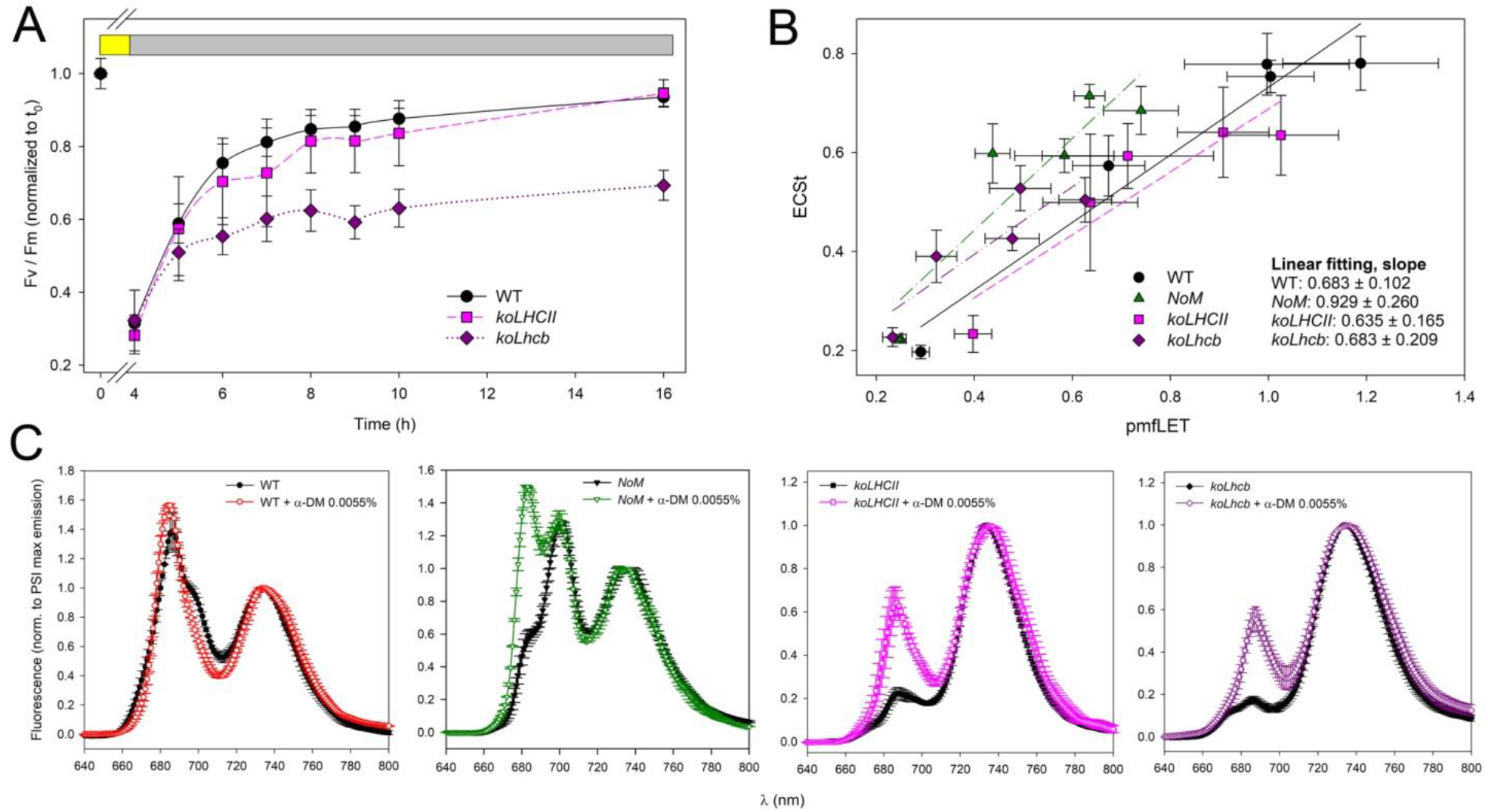
Biophysical characterization of the mechanisms regulating energy transduction reactions in thylakoids. **(A)** PSII repair efficiency. Detached leaves were exposed to EL for 3 hours (wild type, 1100 μmol photons m^−2^ s^−1^; *koLHCII*, 900 μmol photons m^−2^ s^−1^; *koLhcb*, 750 μmol photons m^−2^ s^−1^) at 4°C, to reduce PSII maximal quantum yield (F_v_/F_m_) to about 30% of the initial value. Recovery of F_v_/F_m_ was performed at 15 μmol photons m^−2^ s^−1^, 24°C. *koLhcb* line showed a slower PSII repair kinetic compared to both wild type and *koLHCII*. Values are expressed as mean ± SD, n = 5. **(B)** Relationship between pmfLET and ECSt in wild type and mutant lines upon steady state illumination. All genotypes showed similar amplitude of total ECS as a function of LET, thus suggesting a similar amplitude of CEF. Values are expressed as mean ± SD, n = 4. **(C)** 77K fluorescence emission spectra were recorded on thylakoids from wild type, *NoM, koLHCII* and *koLhcb* plants, before and after treatment with 0.0055% α-DM, severing weak interactions between Chl-binding complexes without solubilizing the membranes. Upon treatment, wild type and *NoM* showed an increased emission by PSII, ascribed to LHCII population disconnected from PSII RC; *koLHCII* and *koLhcb* showed a far higher emission by PSII upon treatment, suggesting the presence of PSII-to-PSI spill-over in these lines. Spectra were normalized to the maximal emission of PSI-LHCI at 733 nm. λ_exc_ = 440 nm. Symbols and error bars show means ± SD, n = 4. See Supplementary Figure S4 for additional details.

We then proceeded to verify whether the different level of thylakoid stacking did affect regulation of LEF *vs*. CEF amplitudes, a crucial mechanism to balance the chloroplast energy budget. To this aim, the kinetic of P700^+^/P700 ratio was monitored in dark-adapted leaves upon far red illumination, when PSII has a far lower turnover rate than PSI. The slow P700 oxidation reflects a high CEF/LEF ratio, because the C3 cycle is inactive in dark-adapted leaves (Joliot & Joliot, 2005) and suggested the capacity to perform CEF in all genotypes. The procedure was repeated at different time intervals, up to 10 minutes upon exposure to actinic light, thus allowing for full rate LEF. When comparing the results from the wild type, *NoM, koLHCII* and *koLhcb*, it clearly appeared that the oxidation rate of P700 increased after sequential periods of priming with light, thus implying that LEF replaced CEF in all genotypes. However, *koLHCII* and *koLhcb* reached lower level of P700^+^/P700 ratio than the wild type, and CEF-to-LEF transition occurred more slowly in *koLhcb* leaves, likely due to their reduced PSII antenna size (Figure 5B).

We then measured Chl fluorescence and the kinetic of electrochromic shift (ECS) on leaves, as a measure of electrons and protons transfer in the photosynthetic apparatus, respectively (Figure 5C). The quantum yield of PSII photochemistry can be used to measure the electron released from PSII (LEF) in leaves under a different light. ECS kinetics can be used to estimate fluxes of protons through the thylakoids, upon steady- state actinic illumination (Cruz et al., 2005): ECS_t_, the total amplitude of the ECS signal upon light-dark transition, measures the light-driven protonmotive force (pmf) across the membrane, while the ECS decay lifetime (gH^+^) assesses the conductivity of the thylakoid ATPase to proton efflux. Under steady-state photosynthetic electron transport, the pmf produced by LEF alone can be estimated by the LEF/gH^+^ ratio, a parameter termed pmfLEF (Avenson et al., 2005). It turned out that the slope of the linear relationship of pmfLEF *vs*. ECS_t_ was proportional to the stoichiometry of electron *vs*. proton transfer. Figure 5C shows that wild type and mutant plants produced approximately the same extent of pmf that can be accounted for by LEF only, thus suggesting a similar CEF amplitude in all genotypes.

Finally, the function of grana stacks, as for light-harvesting, consists in avoiding spill-over of excitons harvested by PSII-LHCII towards PSI-LHCI, which would unbalance the electron transport chain. This is obtained by partitioning PSI-LHCI and PSII-LHCII into two different domains of thylakoids. In order to verify the occurrence of spill-over, we have favored a 77K fluorescence spectroscopy approach. To this aim, we have compared fluorescence emission spectra of thylakoid membranes before and after treatment with very low concentrations of α-DM (0,0055%), which disconnects PSI-LHCI complex from LHCII and/or PSII core complexes in stroma-exposed membranes without solubilization (Caffarri et al., 2014)(Bassi et al., 1985; Caffarri et al., 2014) (Figure 5A). Upon excitation of Chl *a* at 440 nm, the fluorescence-emission spectra of wild-type thylakoids showed three peaks at 686, 695-700 and 735 nm, respectively from Lhcb antenna proteins, PSII core and PSI-LHCI complexes. Emission by PSII can be described with two components, peaking at 684.5 nm for PSII antenna and 692 nm for core complex (Supplementary Figure S4). Upon α-DM addition, the 695 nm shoulder component in 440 nm (Chl *a*) excited spectra was drastically reduced, with a concomitant increase of the 684 nm peak consistent with disruption of excitation energy transfer (EET) from LHCII to PSII and/or PSI. Analysis of spectra from *koLhcb* showed two peaks only, at 689 and 735 nm, from PSII core and PSI-LHCI, respectively. Since the 689 nm peak is weak and is enhanced by α-DM, it can be concluded that PSII core was spilling-over excitation energy to PSI-LHCI and that this EET was severed by α- DM. The analysis of spectra from *koLHCII* confirmed and extended the above conclusions: the untreated thylakoids showed a spectrum very similar to *koLhcb*. Since both components were enhanced (especially the 685 nm peak from LHCs) upon α-DM treatment, we explained the result as the release of quenching by PSI- LHCI on PSII-LHC complexes by α-DM. The case of *NoM* is similar to that of the wild type: besides the broad peak at 735 nm, the major peak occurred at ∼700 nm with a shoulder at 682 nm, a blue-shifted emission component previously ascribed to antennae badly connected to the core (Dall’Osto et al., 2014). Upon α-DM, the 680 nm shoulder was strongly enhanced. This suggests that the α-DM treatment further disconnected the EET from LHCII to PSII core although the effect appeared incomplete when compared with the wild type, as evidenced by the emission at 700 nm from Chl *a* binding core upon excitation at 440 nm.

## Discussion

Grana stacks are the most conspicuous feature of plant chloroplasts, which attracted the attention of Mayer since 1883, when they were first reported (Gunning et al., 2006). The reasons for this interest is that grana are found in land plant, but are absent in algal chloroplast which, despite the obviously common function of chloroplasts through the species, do not have this highly structured multiple membrane layers, but still have differentiation between stacked and unstacked membrane domains (Trissl & Wilhelm, 1993).

The deletion of either the inner layer (monomeric), or the outer layer (trimeric) antenna complexes, or both, had a strong impact on growth rate which was maximal with depletion of monomeric antennas, while the deletion of trimeric LHCII, although causing a moderate decrease in growth, did not affect biomass accumulation as much as it could be predicted by the difference in antenna size (Figure 1A,B). This is particularly evident when comparing the *NoM* with the wild type: since *NoM* strongly over-accumulates LHCII, its PSII antenna size, as determined by fluorescence induction with DCMU, is actually higher than the wild type and yet its growth is decreased by 50% (Figure 2B). We conclude that monomeric LHCs have an essential function in the efficiency of excitation energy transfer from outer antenna to RC of PSII, consistent with the high F_0_ values previously reported for this genotype (Dall’Osto et al., 2014). The higher-than-wild type PSII antenna size in *NoM* is at odd with its reduced growth, implying that factors other than ET to PSII RC are involved in growth retardation. The stronger loss of growth is observed in the absence of both the antenna types. Again, the loss of biomass is far higher than suggested by the apparent reduction antenna size, strengthening the above conclusion. Growth retardation does not appear to be due to enhanced photodamage, because increasing photon fluence by a factor of three (Figure 1B,D) did enhance the growth rate of all the genotypes except for the *koLhcb* plants, whose growth was independent on light intensity. It should be noted that the strongest gain in growth when comparing HL *vs*. LL was obtained in the *koLHCII* plants which retained monomeric LHCs (+80%), suggesting that these complexes are related to such growth promoting factors. The decrease in antenna size of the different phenotypes was associated to an higher PSII/PSI ratio (Figure 2A), a partial compensation for the reduced antenna size of PSII. Indeed, an enhanced PSII *vs*. PSI content and a low F_m_/F_0_ ratio has been observed in the alga *Mantoniella squamata* that lacks grana stacks and thylakoid lateral heterogeneity because of a simplified antenna system composed of a single LHCII like protein (Rhiel & Mörschel, 1993), which is shared by PSI and PSII RCs, leading to spill-over of PSII excitation energy by PSI (Wilhelm et al., 1989).

Based on evidence from basic photosynthetic parameters and the lesson from comparative analysis of thylakoid lateral heterogeneity in *Mantoniella, Pleurochloris, Chlorella* and higher plants (Trissl & Wilhelm, 1993), it could be hypothesized that by knocking out LHC genes we had both decreased the PSII antenna size and reduced the lateral heterogeneity between PSI and PSII, likely by causing de-stacking of thylakoid membranes.

### Thylakoid stacking

Indeed, the analysis of thylakoid stacking confirmed that the growth rate correlated to the extent of thylakoid stacking (Figure 3K). The extent of membrane stacking was decreased by 30% in both *NoM* and *koLHCII* plants, implying that components of both monomeric and trimeric LHC proteins were involved in stabilizing grana partitions. The deletion of both PSII antenna types has a tremendous effect, in that only agranal chloroplasts were detected. The relevance of monomeric vs trimeric LHCs in eliciting grana stacking can be assessed by comparing the *lowLHCII* genotype, containing mostly Lhcb1 with little Lhcb2 gene products, with the *koLHCII*, retaining monomeric complexes. These genotypes had the same overall total LHC antenna level. Since the l*owLHCII* plants had approximatively 18% stacking as compared to the one retaining the three monomeric complexes (*koLHCII*), we conclude that the efficiency of monomeric antennas in causing stacking was 5-6 times higher with respect to trimeric LHCII. It had to be established whether the three inner antennae equally contributed to the stacking, or if this property was specific to one of them. To this aim, we took advantage of the early observation that the *ch1* mutant, in which a lack of Chl *b* de-stabilizes LHCs, retained a substantial level of stacking which closely matched the level found in *koLHCII* (Figure 3K) (Bassi et al., 1985). Loss of Chl *b* prevented accumulation of all LHC proteins, yet CP26 (Lhcb5) was retained (Havaux et al., 2004) because CP26 can binding Chl *a* at most of its Chl *b* binding sites (Croce et al., 2002). When the *ch1 koLhcb5* double mutant was analyzed for thylakoid stacking it showed, indeed, a sharp decrease (Figure 3J), implying that Lhcb5 was a major determinant of grana formation. Last, but not less impressive, was the stacking phenotype of *Lhcb2-only* plants. Lhcb2 is present at low levels in thylakoids as compared to Lhcb1 and yet it was shown to be essential for state 1 - state 2 transitions (Pietrzykowska et al., 2014). State 1 - state 2 transitions cause thylakoid de-stacking upon phosphorylation-dependent migration of LHCII trimers, containing both Lhcb1 and Lhcb2, from grana to stroma membranes (Bassi et al., 1989)(Hepworth et al., 2021). The huge grana stacks, far wider than in the wild type, evidenced in *Lhcb2-only* genotype (Fig. 4B) imply that Lhcb2 is prominent in mediating membrane appression into grana partitions. This was confirmed by the effect of inducing Lhcb2 phosphorylation with PSII light treatment (Figure 4D). Lhcb2 phosphorylation dramatically reduced the number of grana as well as their size (Figure 4B,C,E). We suggest that the grana remaining upon light treatment are promoted by un-phosphorylated LHCII. In a recent study, the formation of dimeric PSII supercomplexes connected by their stromal side was studied as a proxy for thylakoid stacking, by cross-linking and proteomic analysis. Consistent with our finding, both the monomeric and the trimeric antenna were suggested to mediate the interaction between PSII supercomplexes (Albanese et al., 2020). Nevertheless, the identity of the key gene products identified in that study (i.e. Lhcb1 and Lhcb4) does not match our *in vivo* analysis. Although these LHCs might well contribute to stacking, the genetic evidence we show here suggests that the major player in determining stacking and its regulation through phosphorylation are, indeed, the Lhcb5 and the Lhcb2 components of the LHCII trimeric antenna. This difference in attribution might be ascribed to the use of the purified dimeric PSII supercomplex fraction which, although reminiscent of face-to-face stacking, it might represent a PSII population resistant to detergent treatment yet not fully representative of the general PSII organization in grana partitions. Indeed, the same research group has recently reported on alternative organization forms of the face-to-face dimeric complexes (Grinzato et al., 2020) and a recent high resolution study has identified two distinct conformations of the PSII C2S2 complex one of which is consistent with grana stacking being stabilized by interactions between CP26 and a LHCII subunit (Caspy et al., 2021). Alternatively, the differences with respect to our reports might well be ascribed to the *in vivo vs. in vitro* analysis: accessibility of the cross-linkers to individual LHC complexes might not correspond to the strength of interactions they elicit as thylakoid stacking determinants (A. Crepin & Caffarri, 2018).

### Functional significance of the granal stacking

In C3 plants and most green algae, the two PSs are well separated, making the possibility of exciton spilling from PSII antenna in a granal partition to PSI antenna pigments in the stroma-lamellae, rather small (Kirchhoff et al., 2007). This structural arrangement has the advantage of increasing the concentration of PSII, which is kinetically slower, by packing it densely in grana partitions and increasing the cooperativity between PSII RC beyond the obvious level of 2 defined by the dimeric organization of supercomplexes (Lavergne & Joliot, 1991; Lavergne & Trissl, 1995)(Lavergne & Trissl, 1995). The high PSII/PSI stoichiometry ratio and high F_m_/F_0_ ratio are in agreement with such good excitonic separation, which require state 1 - state 2 transitions in order to balance the turnover-rate when light is not equally absorbed by the two PSs (Miller & Lyon, 1985). Moreover, the requirement for the CBB cycle and other metabolic uses of ATP/NADPH from the light phase of photosynthesis, is variable. Indeed, changes in CET *vs*. LET rates have been proposed to depend on state transitions and their effect in modulating the ratio between grana and stroma membranes (Hepworth et al., 2021). Last but not least, PSII is found in its fully functional state in grana partitions, while its repair process upon photoinhibition occurs in stroma membranes, where the machinery for disassembly of PSII core complex is located.

Analysis of these physiological processes in dependence of the extent of grana stacking showed that the efficiency of PSII repair process, i.e. the kinetic of recovery from photoinhibition, was severely affected in *koLhcb* as compared to *koLHCII* (Figure 5A). This effect could be due to the lack of monomeric LHCs in the former genotype or to the lack of grana. We favor the second hypothesis because the *NoM* genotype, although lacking monomeric LHCs, like *koLhcb*, shows recovery kinetic from photoinhibition as in the wild type (Dall’Osto et al., 2020). Instead, we did not observe any major differences in CET *vs*. LET depending on the level of grana stacking, because all genotypes undergo CET to LET transition, although at a different rate (Figure 5B). We interpret the slower transition from CEF to LEF observed in *koLhcb* and, to a lower extent, in *koLHCII*, as caused by the strongly reduced antenna size of this genotype, limiting electron supply and reduction of P700^+^. A strong change in the CET/LET ratio should be reflected in the slope of the electrochromic shift signal *vs*. protonmotive force, which was not observed since all genotypes behave similarly in this respect (Figure 5C).

Lastly, we consider the possibility of excitation energy being spilled over from PSII to PSI upon thylakoid de- stacking. According to previous work with systems showing high spill-over, i.e. the liken *Parmelia sulcata* (Slavov et al., 2013) and the primitive unicellular green alga *Mantoniella squamata* (Trissl & Wilhelm, 1993), a low F_m_/F_0_ ratio associated to the high PSII/PSI stoichiometry is the best indication of ongoing spill-over, as assessed by fast fluorescence spectroscopy at 77K and DAS deconvolution (Slavov et al., 2013). We observed that *koLhcb* has a similarly low F_m_/F_0_ and PSII/PSI ratio as in *Mantoniella* while genotypes retaining significant level of stacking were similar to the wild type, i.e had high F_m_/F_0_ and PSII/PSI ∼1 (Supplementary Table S4). Furthermore, by treating thylakoids with sub-solubilizing concentration of α-DM, which uncouples ET between pigment-proteins in thylakoids (Bassi et al., 1989), we observe a strong up-rise of PSI fluorescence emission bands in the genotypes (*NoM, koLHCII* and *koLhcb*) with a reduced level of stacking, while in the wild type the effect was very small, if any. The case of *koLhcb* is particularly clear, since these thylakoids only contain PSII core and PSI-LHCI complexes, without any PSII antenna (Figure 5A). It should be noted that the canonical method for assessing spill-over consists in showing an increase of P700 antenna size in DCMU-treated samples (Wilhelm et al., 1989). We have attempted this experiment with unconclusive results. Indeed, our mutants, parlicularly koLHCB, have a reduced cross section for photon absorption due to the very nature of the mutations applied: a PSII core complex binds 35 Chl a (Müh & Zouni, 2020) a figure that might be at the limit of detection for optically detected measurement since PSI-LHCI complex binds at least 145 Chls plus several LHCII complexes (Schiphorst et al., 2021). In order to fully verify the hypothesis of spill-over occurring differentially in our grana endowed vs grana depleted genotypes, fast spectroscopy measurements need to be performed and will be the object of future studies. Yet the occurrence of this process in our grana- depleted genotypes is strongly consistent with data, namely the high F_m_/F_0_ ratio of Mantoniella squamata (Rhiel & Mörschel, 1993) and other photosynthetic systems without well defined compatimentation between psi and PSII RC.

We conclude that thylakoid stacking is a characteristic of specific and lowly abundant Lhcb proteins, namely Lhcb5 (CP26) and, chiefly, Lhcb2. Indeed, while CP26 is conserved in algae, Lhcb2 cannot be traced back though the green lineage from land plants to green algae, which, in turn, do not show grana. As for the major physiological functions associated to the evolution of grana, it is a more efficient PSII repair process, possibly due to the compartmentalization of the fully functional PSII separately from the damaged complex under repair and the energy separation of PSI from PSII antenna. Indeed, the evolution of red forms upon land colonization has made the effect of spill-over far more deleterious for an equilibrated energy balance between PSs with respect to algae, whose PSI is depleted in red-forms and is, therefore, less efficient in draining excitation energy from connected antenna beds, according to Boltzmann’s relation.

## Aknowledgements

We thank Pierre Joliot, Francis-André Wollman and Benjamin Bailleul for discussions and suggestions, the team of Electron Microscopy service at the University of Padua-department of Biology Vallisneri for excellent technical assistance. The work was supported by grant RIBA 2017 to R.B. and grant PRIN 2018 to L.D.

## Author contributions

R.B. and L.D. conceived the work and designed the experiments. R.B. performed E.M. analysis. Z.G. carried out the construction of mutants and performed their physiological characterization, together with L.D. and R.C. R.L.G. analyzed fluorescence kinetics and biochemically characterized mutant lines. J.S. developed and provided the CRISPR-Cas9 plasmids system. All authors contributed to writing the manuscript, discussed the results and commented on the manuscript.

## Declaration of Interest

The authors declare no competing interests.

## Materials and Methods

### Plant materials

*Arabidopsis thaliana* mutants *ch1, ch1 koLhcb5, koLhcb3* and *NoM* were obtained as previously described (Dall’Osto et al., 2014; Damkjaer et al., 2009; Havaux et al., 2007). *NoM koLhcb3* was obtained by crossing single mutants and selecting progeny by immunoblotting. *koLHCII, Lhcb2-only, koLhcb* and *lowLHCII* lines were obtained by GE as reported in (Ordon et al., 2017, 2020). Plants were transformed (Zhang et al., 2006) with *Agrobacterium tumefaciens* (strain GV3101). Seedlings were tested for resistance to applications of the antibiotic hygromycin (25 mg l^−1^). For each genotype, independent transformants (T1 generation) were self-fertilized, and absences of proteins were confirmed in the T3 generation by immunotitration.

### Growth conditions

Wild type and mutant genotypes were grown in soil, in a phytotron for 6 weeks at either 150 or 350 μmol photons m^−2^ s^−1^, 23°C, 70% relative humidity, 8/16 h of day/night. All biochemical and physiological analyses were performed on plants prior to the onset of flowering. The growth of plants was determined by measuring the rosette fresh weight at the end of the growth cycle.

### Membrane isolation

Stacked thylakoid membranes were isolated as previously described (Casazza et al., 2001).

### Pigment analysis

Pigments were extracted from leaf discs with 85% acetone buffered with Na_2_CO_3_, then separated and quantified by HPLC (Jasco Extrema LC-4000) as in (Gilmore & Yamamoto, 1991).

### Electrophoresis and immunoblotting

SDS-PAGE analysis of thylakoid proteins was performed using the Tris-Tricine buffer system (Schägger & von Jagow, 1987). For immunotitration (Towbin et al., 1979), proteins were detected with alkaline phosphatase-conjugated antibody (Sigma-Aldrich A3687). Primary antibodies used were: α-PsbB/CP47 (AS04 038), α-Lhcb1 (AS01 004), α-Lhcb2 (AS01 003), α-P-Lhcb1 (AS13 2704), α-P-Lhcb2 (AS13 2705) from Agrisera. Signal amplitude was quantified using the GelPro 3.2 software (Bio-Rad). Non-denaturing Deriphat-PAGE was performed as in (Guardini et al., 2020). SDS-PAGE analysis on complexes eluted from non-denaturing gel was performed using the Laemmli system as reported in (Aurélie Crepin et al., 2020).

### Spectroscopy

Absorption spectra were recorded at RT using a Jasco V-550 spectrophotometer. Leaf pigment content was calculated from spectra of acetonic extracts (Croce et al., 2002). Absorption measurements on complexes eluted from non-denaturing gel were performed in 5% glycerol, 10 mM HEPES pH 7.5, α-DM 0.004%. 77K fluorescence emission spectra (λ_exc_ 440 nm, ∼0.2 µg Chl mL^-1^ in 50% glycerol, 10 mM HEPES, pH 7.5) were recorded using a Jobin-Yvon Fluoromax-3 spectrofluorometer (Dall’Osto et al., 2017).

### Analysis of Chl fluorescence

Photosynthetic parameters (Baker, 2008) were measured on leaves at RT with a PAM 101 fluorimeter (Heinz-Walz). Changes in the redox state of P700 and CEF to LEF transition were measured as in (Dal Corso et al., 2008) with a DUAL-PAM-100 equipment (Walz, GmbH). Variable fluorescence was measured in a home-built Chl fluorimeter, and induced by green light (Rappaport et al., 2007) (10 μmol photons m^−2^ s^−1^) in dark adapted leaves, infiltrated with DCMU 50 μM. Steady state, light- induced pmf was estimated from changes in absorbance associated with the ECS at 520 nm (Cruz et al., 2005; Livingston et al., 2010), using a LED spectrophotometer (JTS10; Biologic Science Instruments).

### Electron Microscopy and image analysis

Transmission electron microscopy on leaf fragments was conducted using a FEI Tecnai T12 electron microscope operating at 100 kV accelerating voltage. Leaf fragments were fixed in 3% glutaraldehyde in 0.1 M cacodylate buffer pH 6.9. Analyses on EM images were conducted with ImageJ software (Schneider et al., 2012).

### LHCII phosphorylation induction

Leaves were dark adapted for 60 minutes prior to exposure to 200 μmol photons m^-2^ s^-1^ PSII light (orange) for 60 minutes at 22°C.

### Statistics

Statistical analyses were performed in SigmaPlot using One-way analysis of variance (ANOVA), means were separated with Tukey’s post hoc test at a significant level of P < 0.05 (see the figure legends for details). Error bars represent the standard deviation.

### Data availability

Sequence data from this article can be found in the *Arabidopsis* Genome Initiative or GenBank/EMBL databases under accession numbers At1g29920 *(Lhcb1.1*), At1g29910 (*Lhcb1.2*), At1g29930 (*Lhcb1.3*), At2g34430 (*Lhcb1.4*), At2g34420 (*Lhcb1.5*), At2g05100 (*Lhcb2.1*), At2g05070 (*Lhcb2.2*), At3g27690 (*Lhcb2.3*), At5g54270 (*Lhcb3*), At5g01530 (*Lhcb4.1*), At3g08940 (*Lhcb4.2*), At4g10340 (*Lhcb5*), At1g44446 (*cao*). The KO lines used in the work were obtained from the NASC under the stock numbers N376476 (*koLhcb4.1*), N877954 (*koLhcb4.2*), N514869 (*koLhcb5*), N520342 (*koLhcb3*), N524295 (*ch1*).

## Notes

### Competing Interest Statement

The authors have declared no competing interest.

## References

Albanese, P., Tamara, S., Saracco, G., Scheltema, R. A., & Pagliano, C. (2020). How paired PSII–LHCII supercomplexes mediate the stacking of plant thylakoid membranes unveiled by structural mass-spectrometry. Nature Communications, 11(1). https://doi.org/10.1038/s41467-020-15184-1

Allred, D. R., & Staehelin, L. A. (1985). Lateral distribution of the cytochrome b6/f and coupling factor ATP-synthetase complexes of chloroplast thylakoid membranes. Plant Physiol., 78, 199–202.

Anderson, J. M., & Melis, A. (1983). Localization of different photosystems in seperate regions of chloroplast membranes. Proc.Natl.Acad.Sci.USA, 80, 745–749.

Argyroudi-Akoyunoglou, J. H., Feleki, Z., & Akoyunoglou, G. (1971). Formation of two chlorophyll-protein complexes during greening of etiolated bean leaves. Biochemical and Biophysical Research Communications, 45(3), 606–614. https://doi.org/10.1016/0006-291X(71)90460-8

Armond, P. A., & Arntzen, C. J. (1977). Localization and Characterization of Photosystem II in Grana and Stroma Lamellae. Plant Physiology, 59(3), 398–404. https://doi.org/10.1104/pp.59.3.398

Armond, P. A., Arntzen, C. J., Briantais, J. M., & Vernotte, C. (1976). Differentiation of chloroplast lamellae. Light harvesting efficiency and grana development. Archives of Biochemistry and Biophysics, 175(1), 54–63. https://doi.org/10.1016/0003-9861(76)90484-7

Armond, P. A., Staehelin, L. A., & Arntzen, C. J. (1977). Spatial relationship of photosysten I, photosystem II, and the light-harvesting complex in chloroplast membranes. J.Cell Biol., 73, 400–418.

Avenson, T. J., Cruz, J. A., Kanazawa, A., & Kramer, D. M. (2005). Regulating the proton budget of higher plant photosynthesis. Proceedings of the National Academy of Sciences of the United States of America, 102(27), 9709–9713. https://doi.org/10.1073/pnas.0503952102

Baker, N. R. (2008). Chlorophyll Fluorescence: A Probe of Photosynthesis In Vivo. Annual Review of Plant Biology, 59(1), 89–113. https://doi.org/10.1146/annurev.arplant.59.032607.092759

Bassi, R., Ghiretti Magaldi, A., Tognon, G., Giacometti, G. M., & Miller, K. R. (1989). Two-dimensional crystals of the photosystem II reaction center complex from higher plants. European Journal of Cell Biology, 50(1), 84–93. https://europepmc.org/article/med/2693092

Bassi, R., Marquardt, J., & Lavergne, J. (1995). Biochemical and fuctional properties of photosystem II in agranal membranes from maize mesophyll and bundle sheath chloroplasts. Eur.J.Biochem., 233, 709–719.

Bassi, R., & Simpson, D. J. (1986). Differential expression of LHCII genes in mesophyll and bundle sheath cells of maize. Carlsberg Research Communications, 51(6), 363–370. https://doi.org/10.1007/BF02907312

Bassi, R, Hinz, U., & Barbato, R. (1985). The role of the light harvesting complex and photosystem II in thylakoid stacking in the chlorina-f2 barley mutant. Carlsberg Res.Commun., 50_ _, 347–367.

Belgio, E., Kapitonova, E., Chmeliov, J., Duffy, C. D. P., Ungerer, P., Valkunas, L., & Ruban, A. V. (2014). Economic photoprotection in photosystem II that retains a complete light-harvesting system with slow energy traps. Nature Communications, 5, 1–8. https://doi.org/10.1038/ncomms5433

Caffarri, S., Tibiletti, T., Jennings, R., & Santabarbara, S. (2014). A Comparison Between Plant Photosystem I and Photosystem II Architecture and Functioning. Current Protein & Peptide Science, 15(4), 296–331. https://doi.org/10.2174/1389203715666140327102218

Campoli, C., Caffarri, S., Svensson, J. T., Bassi, R., Stanca, A. M., Cattivelli, L., & Crosatti, C. (2009). Parallel pigment and transcriptomic analysis of four barley Albina and Xantha mutants reveals the complex network of the chloroplast-dependent metabolism. Plant Molecular Biology, 71(1–2), 173–191.

Casazza, A. P., Tarantino, D., & Soave, C. (2001). Preparation and functional characterization of thylakoids from Arabidopsis thaliana. Photosynth.Res., 68(0166-8595 (Print)), 175–180. https://doi.org/10.1023/A:1011818021875

Caspy, I., Fadeeva, M., Mazor, Y., & Nelson, N. (2021). Structure of Dunaliella Photosystem II reveals conformational flexibility of stacked and unstacked supercomplexes. https://doi.org/10.1101/2021.11.29.470333

Crepin, A., & Caffarri, S. (2018). Functions and evolution of Lhcb isoforms composing LHCII, the major light harvesting complex of Photosystem II of green eukaryotic organisms. Current Protein & Peptide Science, 19. https://doi.org/10.2174/1389203719666180222101534

Crepin, Aurélie, Kučerová, Z., Kosta, A., Durand, E., & Caffarri, S. (2020). Isolation and characterization of a large photosystem I–light-harvesting complex II supercomplex with an additional Lhca1–a4 dimer in Arabidopsis. Plant Journal, 102(2), 398–409. https://doi.org/10.1111/tpj.14634

Croce, R., Canino, G., Ros, F., & Bassi, R. (2002). Chromophore organization in the higher-plant photosystem II antenna protein CP26. Biochemistry, 41(23), 7334–7343. https://doi.org/10.1021/bi0257437

Cruz, J. A., Avenson, T. J., Kanazawa, A., Takizawa, K., Edwards, G. E., & Kramer, D. M. (2005). Plasticity in light reactions of photosynthesis for energy production and photoprotection. Journal of Experimental Botany, 56(411), 395–406. https://doi.org/10.1093/jxb/eri022

Dal Corso, G., Pesaresi, P., Masiero, S., Aseeva, E., Schünemann, D., Finazzi, G., Joliot, P., Barbato, R., & Leister, D. (2008). A Complex Containing PGRL1 and PGR5 Is Involved in the Switch between Linear and Cyclic Electron Flow in Arabidopsis. Cell, 132(2), 273–285. https://doi.org/10.1016/j.cell.2007.12.028

Dall’Osto, L., Cazzaniga, S., Bressan, M., Paleček, D., Židek, K., Niyogi, K. K., Fleming, G. R., Zigmantas, D., & Bassi, R. (2017). Two mechanisms for dissipation of excess light in monomeric and trimeric light-harvesting complexes. Nature Plants, 3(April), 17033. https://doi.org/10.1038/nplants.2017.33

Dall’Osto, L., Cazzaniga, S., Zappone, D., & Bassi, R. (2020). Monomeric light harvesting complexes enhance excitation energy transfer from LHCII to PSII and control their lateral spacing in thylakoids. Biochimica et Biophysica Acta - Bioenergetics, 1861(4), 0–1. https://doi.org/10.1016/j.bbabio.2019.06.007

Dall’Osto, L., Ünlü, C., Cazzaniga, S., & Van Amerongen, H. (2014). Disturbed excitation energy transfer in Arabidopsis thaliana mutants lacking minor antenna complexes of photosystem II. BBA - Bioenergetics, 1837(12), 1981–1988. https://doi.org/10.1016/j.bbabio.2014.09.011

Damkjaer, J., Kereiche, S., Johnson, M. P., Kovacs, L., a.z Kiss, Boekema, E. J., Ruban, A. V, Horton, P., & Jansson, S. (2009). The Photosystem II light-harvesting protein Lhcb3 affects the macrostructure of Photosystem II and the rate of state transitions in Arabidopsis. Plant Cell, 21, 3245–3256.

Gilmore, A. M., & Yamamoto, H. Y. (1991). Zeaxanthin Formation and Energy-Dependent Fluorescence Quenching in Pea Chloroplasts under Artificially Mediated Linear and Cyclic Electron Transport1. Plant Physiology, 96(2), 635–643. https://doi.org/10.1104/pp.96.2.635

Granick, S., & Porter, K. R. (1947). THE STRUCTURE OF THE SPINACH CHLOROPLAST AS INTERPRETED WITH THE ELECTRON MICROSCOPE. American Journal of Botany, 34(10), 545–550. https://doi.org/10.1002/J.1537-2197.1947.TB13028.X

Grinzato, A., Albanese, P., Marotta, R., Swuec, P., Saracco, G., Bolognesi, M., Zanotti, G., & Pagliano, C. (2020). High-Light versus Low-Light: Effects on Paired Photosystem II Supercomplex Structural Rearrangement in Pea Plants. International Journal of Molecular Sciences, 21(22), 8643. https://doi.org/10.3390/ijms21228643

Guardini, Z., Bressan, M., Caferri, R., Bassi, R., & Dall’Osto, L. (2020). Identification of a pigment cluster catalysing fast photoprotective quenching response in CP29. Nature Plants, 6(3), 303–313. https://doi.org/10.1038/s41477-020-0612-8

Gunning, B., Koenig, F., & Govindjee. (2006). A Dedication to Pioneers of Research on Chloroplast Structure. In R. R. Wise & J. K. Hoober (Eds.), The structure and function of plastids (1st ed., pp. xxiii–xxxi). Springer.

Havaux, M., Dall’Osto, L., & Bassi, R. (2007). Zeaxanthin has enhanced antioxidant capacity with respect to all other xanthophylls in Arabidopsis leaves and functions independent of binding to PSII antennae. Plant Physiol, 145(0032-0889 (Print)), 1506–1520.

Havaux, M., Dall’Osto, L., Cuiné, S., Giuliano, G., & Bassi, R. (2004). The Effect of Zeaxanthin As the only Xanthophyll on the Structure and Function of the Photosynthetic Apparatus in Arabidopsis thaliana. Journal of Biological Chemistry, 279(14), 13878–13888. https://doi.org/10.1074/jbc.M311154200

Heinz, S., Rast, A., Shao, L., Gutu, A., Gügel, I. L., Heyno, E., Labs, M., Rengstl, B., Viola, S., Nowaczyk, M. M., Leister, D., & Nickelsen, J. (2016). Thylakoid membrane architecture in Synechocystis depends on CurT, a homolog of the granal CURVATURE THYLAKOID1 proteins. Plant Cell, 28(9), 2238–2260. https://doi.org/10.1105/tpc.16.00491

Heitz, E. (1936). Untersuchungen Über den Bau der Plastiden. Planta 1936 26:1, 26(1), 134–163. https://doi.org/10.1007/BF01913844

Hepworth, C., Wood, W. H. J., Emrich-Mills, T. Z., Proctor, M. S., Casson, S., & Johnson, M. P. (2021). Dynamic thylakoid stacking and state transitions work synergistically to avoid acceptor-side limitation of photosystem I. Nature Plants, 7(1), 87–98. https://doi.org/10.1038/s41477-020-00828-3

Iwai, M., Roth, M. S., & Niyogi, K. K. (2018). Subdiffraction-resolution live-cell imaging for visualizing thylakoid membranes. Plant Journal, 96(1), 233–243. https://doi.org/10.1111/tpj.14021

Joliot, P., & Joliot, A. (2005). Quantification of cyclic and linear flows in plants. Proc.Natl.Acad.Sci.U.S.A, 102(0027-8424 (Print)), 4913–4918.

Kim, E. H., Li, X. P., Razeghifard, R., Anderson, J. M., Niyogi, K. K., Pogson, B. J., & Chow, W. S. (2009). The multiple roles of light-harvesting chlorophyll a/b-protein complexes define structure and optimize function of Arabidopsis chloroplasts: A study using two chlorophyll b-less mutants. Biochimica et Biophysica Acta - Bioenergetics, 1787(8), 973–984. https://doi.org/10.1016/j.bbabio.2009.04.009

Kirchhoff, H. (2019). Chloroplast ultrastructure in plants. New Phytologist, 223(2), 565–574. https://doi.org/10.1111/NPH.15730

Kirchhoff, H., Haase, W., Haferkamp, S., Schott, T., Borinski, M., Kubitscheck, U., & Rögner, M. (2007). Structural and functional self-organization of Photosystem II in grana thylakoids. Biochimica et Biophysica Acta (BBA) - Bioenergetics, 1767(9), 1180–1188. https://doi.org/10.1016/J.BBABIO.2007.05.009

Kirchhoff, H., Hall, C., Wood, M., Herbstová, M., Tsabari, O., Nevo, R., Charuvi, D., Shimoni, E., & Reich, Z. (2011). Dynamic control of protein diffusion within the granal thylakoid lumen. Proceedings of the National Academy of Sciences of the United States of America, 108(50), 20248–20253. https://doi.org/10.1073/pnas.1104141109

Koochak, H., Puthiyaveetil, S., Mullendore, D. L., Li, M., & Kirchhoff, H. (2019). The structural and functional domains of plant thylakoid membranes. Plant Journal, 97(3), 412–429. https://doi.org/10.1111/tpj.14127

Kramer, D. M., & Evans, J. R. (2011). The importance of energy balance in improving photosynthetic productivity. Plant Physiology, 155(1), 70–78. https://doi.org/10.1104/pp.110.166652

Kyle, D. J., Staehelin, L. A., & Arntzen, C. J. (1983). Lateral mobility of the light-harvesting complex in chloroplast membranes controls excitation energy distribution in higher plants. Arch.Biochem.Biophys., 222, 527–541.

Lavergne, J., & Joliot, P. (1991). Restricted Diffusion in Photosynthetic Membranes. Trends Biochem.Sci., 16, 129–134.

Lavergne, J., & Trissl, H. W. (1995). Theory of fluorescence induction in photosystem II: derivation of analytical expressions in a model including exciton-radical-pair equilibrium and restricted energy transfer between photosynthetic units. Biophysical Journal, 68(6), 2474–2492. https://doi.org/10.1016/S0006-3495(95)80429-7

Livingston, A. K., Cruz, J. A., Kohzuma, K., Dhingra, A., & Kramer, D. M. (2010). An arabidopsis mutant with high cyclic electron flow around photosystem i (hcef) involving the nadphdehydrogenase complex. Plant Cell, 22(1), 221–233. https://doi.org/10.1105/tpc.109.071084

Malzahn, A., Lowder, L., & Qi, Y. (2017). Plant genome editing with TALEN and CRISPR. Cell and Bioscience, 7(1), 1–18. https://doi.org/10.1186/s13578-017-0148-4

Margulies, M. M. (1966). Effect of Chloramphenicol on Formation of Chloroplast Structure and Protein During Greening of Etiolated Leaves of Phaseolus vulgaris. Plant Physiology, 41(6), 992–1003. https://doi.org/10.1104/pp.41.6.992

Mattoo, A. K., Giardi, M.-T., Raskind, A., & Edelman, M. (1999). Dynamic metabolism of photosystem II reaction center proteins and pigments. Physiologia Plantarum, 107(4), 454–461. https://doi.org/10.1034/J.1399-3054.1999.100412.X

Mattoo, A. K., Marder, J. B., & Edelman, M. (1989). Dynamics of the photosystem II reaction center. Cell, 56, 241–246.

Miller, K. R., & Lyon, M. K. (1985). Do we really know why chloroplast membranes stack? Trends in Biochemical Sciences, 10(6), 219–222. https://doi.org/10.1016/0968-0004(85)90132-X

Müh, F., & Zouni, A. (2020). Structural basis of light-harvesting in the photosystem II core complex. Protein Science, 29(5), 1090–1119. https://doi.org/10.1002/pro.3841

Mullineaux, C. W. (2005). Function and evolution of grana. Trends in Plant Science, 10(11), 521–525. https://doi.org/10.1016/j.tplants.2005.09.001

Mullineaux, C. W., & Emlyn-Jones, D. (2005). State transitions: An example of acclimation to low-light stress. Journal of Experimental Botany, 56(411), 389–393. https://doi.org/10.1093/jxb/eri064

Murata, N. (1969). Control of excitation transfer in photosynthesis. I. Light-induced change of chlorophyll a fluorescence in Porphyridium cruentum. Biochim.Biophys.Acta, 172, 242–251.

Olive, J., Vallon, O., Wollman, F. A., Recouvreur, M., & Bennoun, P. (1986). Studies on the Cytochrome B6/F Complex .2. Localization of the Complex in the Thylakoid Membranes from Spinach and Chlamydomonas-Reinhardtii by Immunocytochemistry and Freeze-Fracture Analysis of B6/F Mutants. Biochimica et Biophysica Acta, 851(2), 239–248.

Ordon, J., Bressan, M., Kretschmer, C., Dall’Osto, L., Marillonnet, S., Bassi, R., & Stuttmann, J. (2020). Optimized Cas9 expression systems for highly efficient Arabidopsis genome editing facilitate isolation of complex alleles in a single generation. Functional and Integrative Genomics, 20(1), 151–162. https://doi.org/10.1007/s10142-019-00665-4

Ordon, J., Gantner, J., Kemna, J., Schwalgun, L., Reschke, M., Streubel, J., Boch, J., & Stuttmann, J. (2017). Generation of chromosomal deletions in dicotyledonous plants employing a user-friendly genome editing toolkit. Plant Journal, 89(1), 155–168. https://doi.org/10.1111/tpj.13319

Paolillo, D. J. J., & Reighard, J. A. (2011). ON THE RELATIONSHIP BETWEEN MATURE STRUCTURE AND ONTOGENY IN THE GRANA OF CHLOROPLASTS. https://Doi.Org/10.1139/B67-083, 45(6), 773–782. https://doi.org/10.1139/B67-083

Pietrzykowska, M., Suorsa, M., Semchonok, D. A., Tikkanen, M., Boekema, E. J., Aro, E. M., & Jansson, S. (2014). The light-harvesting chlorophyll a/b binding proteins Lhcb1 and Lhcb2 play complementary roles during state transitions in Arabidopsis. Plant Cell, 26(9), 3646–3660. https://doi.org/10.1105/tpc.114.127373

Pribil, M., Labs, M., & Leister, D. (2014). Structure and dynamics of thylakoids in land plants. In Journal of Experimental Botany (Vol. 65, Issue 8, pp. 1955–1972). J Exp Bot. https://doi.org/10.1093/jxb/eru090

Rappaport, F., Béal, D., Joliot, A., & Joliot, P. (2007). On the advantages of using green light to study fluorescence yield changes in leaves. 1767, 56–65. https://doi.org/10.1016/j.bbabio.2006.10.002

Rhiel, E., & Mörschel, E. (1993). The atypical chlorophyll a/b/c light-harvesting complex of Mantoniella squamata: molecular cloning and sequence analysis. Molecular and General Genetics MGG 1993 240:3, 240(3), 403–413. https://doi.org/10.1007/BF00280392

Schägger, H., & von Jagow, G. (1987). Tricine-sodium dodecyl sulfate-polyacrylamide gel electrophoresis for the separation of proteins in the range from 1 to 100 kDa. Anal.Biochem., 166(2), 368–379. https://doi.org/10.1016/0003-2697(87)90587-2

Schiphorst, C., Achterberg, L., Go, R., Koehorst, R., Bassi, R., Amerongen, H. Van, Osto, L. D., Wientjes, E., & Biotecnologie, D. (2021). The role of light-harvesting complex I in excitation energy transfer from LHCII to photosystem I in Arabidopsis. 1–12.

Schiphorst, C., & Bassi, R. (2020). Chlorophyll-Xanthophyll Antenna Complexes: In Between Light Harvesting and Energy Dissipation. 27–55. https://doi.org/10.1007/978-3-030-33397-3_3

Schneider, C. A., Rasband, W. S., & Eliceiri, K. W. (2012). NIH Image to ImageJ: 25 years of image analysis. https://doi.org/10.1038/nmeth.2089

Simpson, D. J. (1982). Freeze-fracture studies on barley plastid membranes V.viridis-n 34, a photosystem I mutant. Carlsberg Research Communications 1982 47:4, 47(4), 215–225. https://doi.org/10.1007/BF02907873

Simpson, D. J., & Robinson, S. P. (1984). Freeze-Fracture Ultrastructure of Thylakoid Membranes in Chloroplasts from Manganese-Deficient Plants. Plant Physiology, 74(3), 735–741. https://doi.org/10.1104/PP.74.3.735

Simpson, D. J., Vallon, O., & von Wettstein, D. (1989). Freeze-fracture studies on barley plastid membranes: VIII. In viridis-115, a mutant completely lacking Photosystem II, oxygen evolution enhancer 1 (OEE1) and the α-subunit of cytochrome b-559 accumulate in appressed thylakoids. Biochimica et Biophysica Acta (BBA) - Bioenergetics, 975(1), 164–174. https://doi.org/10.1016/S0005-2728(89)80215-4

Slavov, C., Reus, M., & Holzwarth, A. R. (2013). Two Different Mechanisms Cooperate In The Desiccation-Induced Excited State Quenching In Parmelia Lichen. Journal of Physical Chemistry B, 117(38), 11326–11336. https://doi.org/10.1021/JP402881F

Standfuss, R., van Scheltinga, A. C. T., Lamborghini, M., & Kuhlbrandt, W. (2005). Mechanisms of photoprotection and nonphotochemical quenching in pea light-harvesting complex at 2.5A resolution. Embo Journal, 24(5), 919–928.

Towbin, H., Staehelin, T., & Gordon, J. (1979). Electrophoretic transfer of proteins from polyacrylamide gels to nitrocellulose sheets: Procedure and some applications. Proceedings of the National Academy of Sciences, 76(9), 4350–4354. https://doi.org/10.1073/pnas.76.9.4350

Trissl, H. W., & Wilhelm, C. (1993). Why do thylakoid membranes from higher plants form grana stacks? Trends in Biochemical Sciences, 18(11), 415–419. https://doi.org/10.1016/0968-0004(93)90136-B

Vallon, O., Bulte, L., Dainese, P., Olive, J., Bassi, R., & Wollman, F. A. (1991). Lateral redistribution of cytochrome b6/f complexes along thylakoid membranes upon state transitions. Proc.Natl.Acad.Sci.U.S.A, 88(0027-8424 SB-IM), 8262–8266.

Wietrzynski, W., Schaffer, M., Tegunov, D., Albert, S., Kanazawa, A., Plitzko, J. M., Baumeister, W., & Engel, B. D. (2020). Charting the native architecture of Chlamydomonas thylakoid membranes with single-molecule precision. ELife, 9, 1–18. https://doi.org/10.7554/eLife.53740

Wilhelm, C., Krämer, P., & Lenartz-Weiler, I. (1989). The energy distribution between the photosystems and light-induced changes in the stoichiometry of system I and II reaction centers in the chlorophyll b-containing alga Mantoniella squamata (Prasinophyceae). Photosynthesis Research 1989 20:3, 20(3), 221–233. https://doi.org/10.1007/BF00034066

Zhang, X., Henriques, R., Lin, S.-S., Niu, Q.-W., Chua, N.-H., Niu, W.-W., Chua, N.-H., Niu, Q.-W., & Chua, N.-H. H. N.-H. (2006). Agrobacterium-mediated transformation of Arabidopsis thaliana using the floral dip method. Nature Protocols, 1(2), 641–646. https://doi.org/10.1038/nprot.2006.97

