## Supplementary Information for "Thylakoid grana stacking revealed by multiplex genome editing of LHCII encoding genes"

### Supplementary Results

#### Supplementary Figure S1

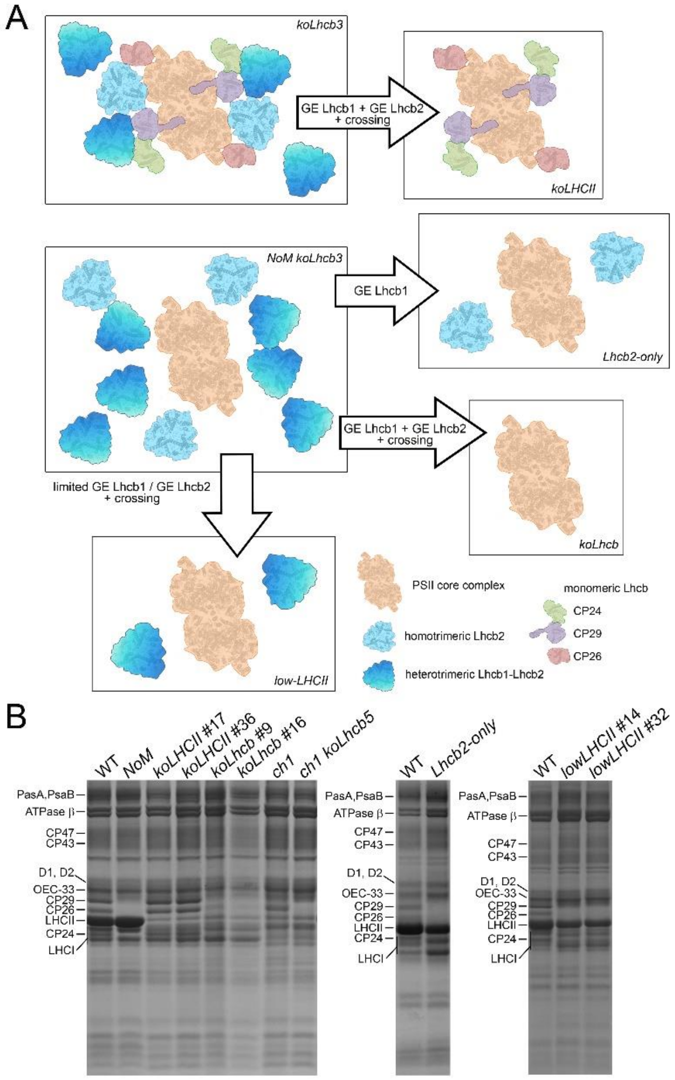


**Supplementary Figure S1. Schematic diagram and protein composition of *Arabidopsis* LHC mutants produced in the present work. (A)** The *NoM* line was obtained as previously reported (Dall'Osto et al. 2017), and a T-DNA insertion in At5g54270 (Lhcb3; NASC N520342) was further introgressed. Specific sgRNA were selected and used for the deletion of either *Lhcb1* or *Lhcb2* genes (Ordon et al. 2020), in both *koLhcb3* and *NoM koLhcb3* genetic backgrounds. *koLHCII* and *koLhcb* lines were obtained by crossing *koLhcb1* x *koLhcb2* lines and selecting progeny by immunoblotting, while *Lhcb2-only* lines were obtained by deleting *Lhcb1* genes in *NoM koLhcb3* plants. *low-LHCII* lines were produced by crossing *NoM koLhcb3* lines which underwent limited genome editing on *Lhcb1* and *Lhcb2* loci, then selecting in the progeny plants retaining ~30% of LHC content than parental genotype. **(B)** SDS–PAGE analysis of wild type (WT) and LHC mutant thylakoid proteins, performed with the Tris-Tricine buffer system. Selected apoprotein bands are marked.

**Supplementary Figure S2**

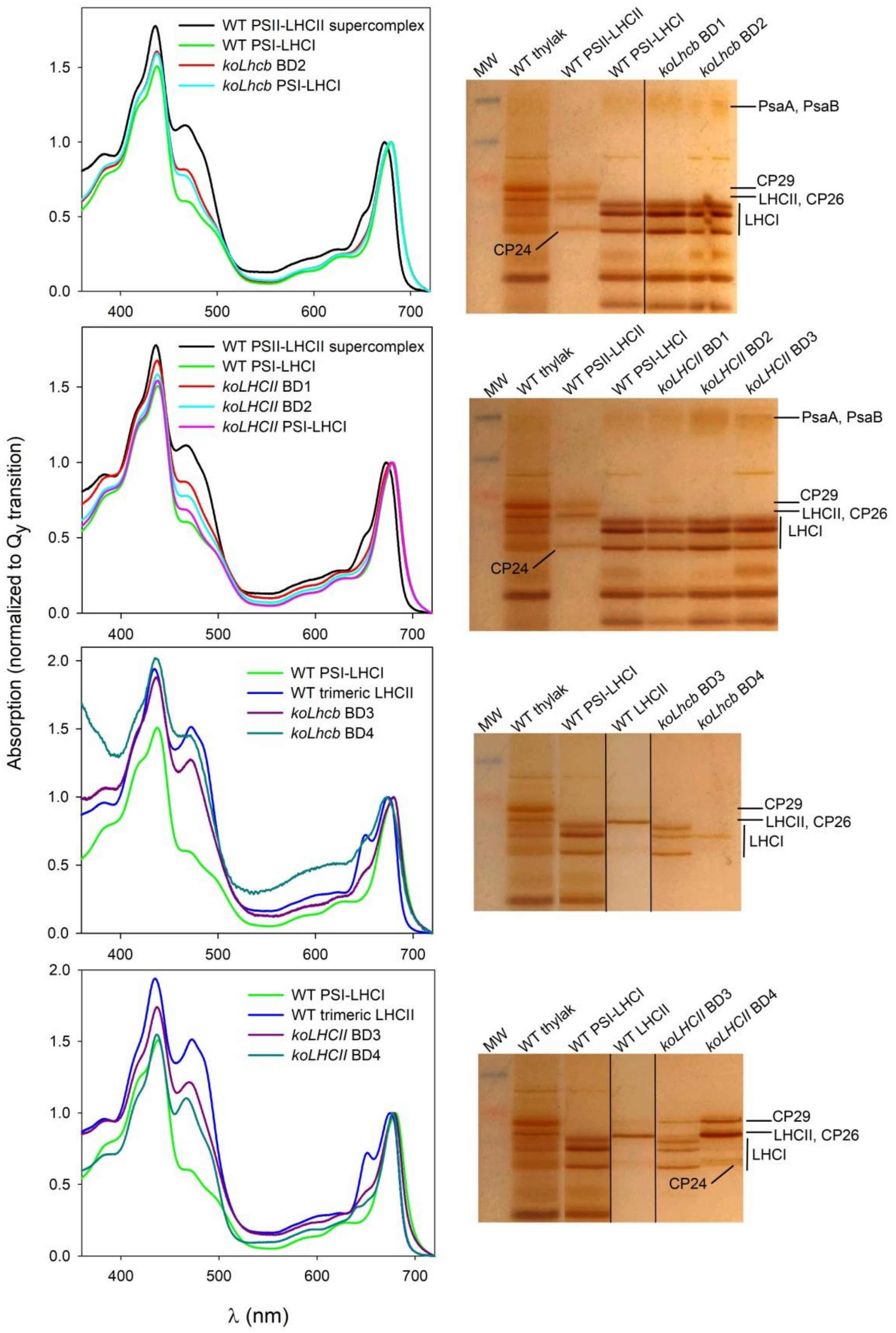


**Supplementary Figure S2. Insights on the composition of pigment-binding complexes separated by non-denaturing Deriphat-PAGE (see Figure 2A).** Chl-binding complexes were extracted from acrylamide matrix and further characterized by absorption spectroscopy at RT (*left column*). Moreover, protein composition of selected bands was determined by SDS-PAGE and silver staining (*right column*). Fractions BD1 and BD2 contained PSI-LHCI supercomplexes with increasing

LHC content, as determined by (i) the red–tail of the $Q_y$ transition, and (ii) the lower Chl *a/b* ratio with respect to the PSI-LHCI supercomplex from WT thylakoids. In all these bands, major subunits identified were PsaA/PsaB (PSI core complex) and LHCI (PSI antennae). BD3 and BD4 from *koLhcb* contained LHCI, likely dimeric in BD3 (which indeed showed an electrophoretic mobility intermediate between trimeric LHCII and monomeric Lhcb), while monomeric in BD4. BD3 from *koLHCII* was enriched in LHCI subunits, but also contained dimeric states of Lhcb complex (CP29/CP26). In BD4 from *koLHCII*, the relative abundance of monomeric LHCI was reduced than BD3, while the monomeric Lhcb CP29, CP26 and CP24 were the major components.

**Supplementary Figure S3**

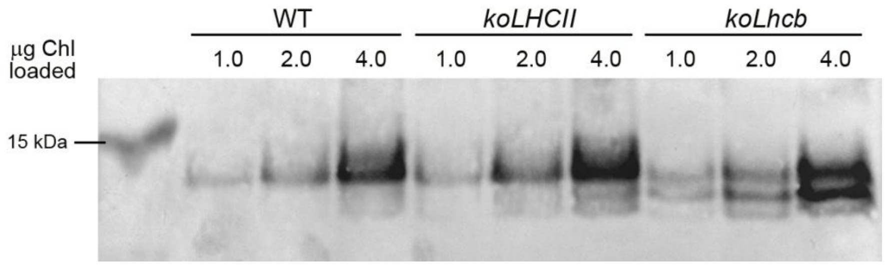


**Supplementary Figure S3. Analysis of CURT1, the factor controlling curvature of thylakoid membranes at the margins and grana shaping.** Immunoblotting used for the quantification of the CURT1A subunits in the WT and mutant thylakoids. Proteins corresponding to 1.0, 2.0 and 4.0 µg of Chls were loaded for each sample. All samples were loaded on the same SDS-PAGE slab gel.

### Supplementary Figure S4

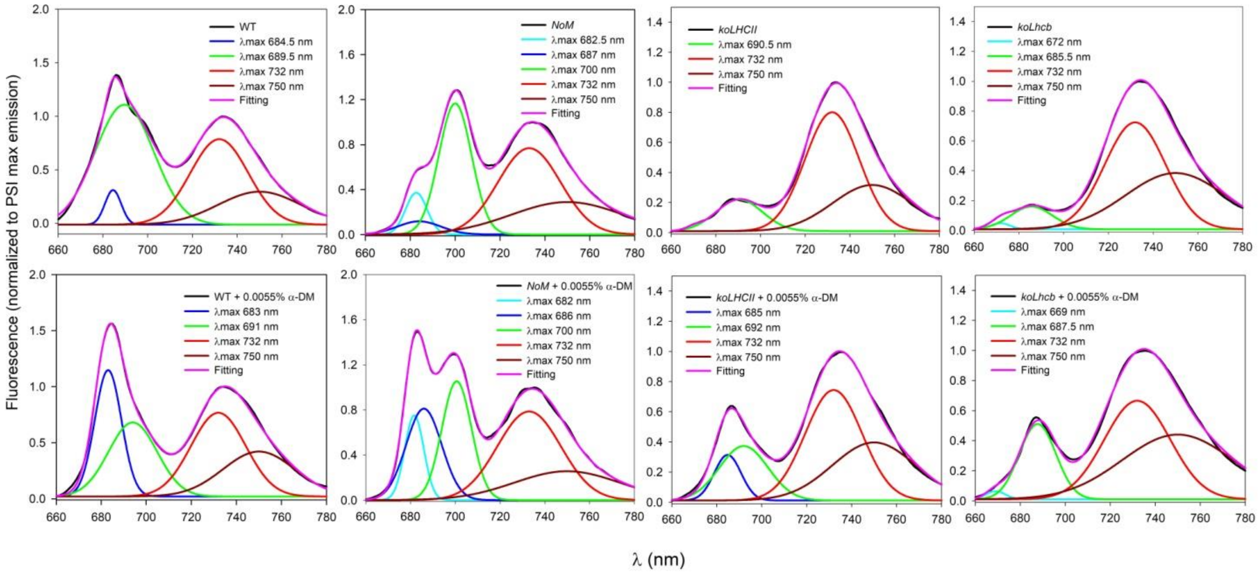


| | | | | | |
|---|---|---|---|---|---|
| **WT** | | | **peak 683-684 nm** | **peak 689-691 nm** | **peaks 732+750 nm** |
| | ctrl | | 7.3 ± 0.6 | 92.2 ± 0.8 | 100.0 ± 1.7 |
| | + α-DM 0.0055% | | 43.3 ± 0.6 | 46.1 ± 1.2 | 100.0 ± 1.6 |
| ***NoM*** | | **peak 682 nm** | **peak 686-687 nm** | **peak 700 nm** | **peaks 732+750 nm** |
| | ctrl | 9.8 ± 2.9 | 6.2 ± 6.9 | 47.4 ± 4.7 | 100.0 ± 1.5 |
| | + α-DM 0.0055% | 16.6 ± 1.3 | 37.4 ± 2.1 | 40.5 ± 0.9 | 100.0 ± 1.1 |
| ***koLHCII*** | | | **peak 685 nm** | **peak 690-692 nm** | **peaks 732+750 nm** |
| | ctrl | | - | 14.5 ± 2.1 | 100.0 ± 1.0 |
| | + α-DM 0.0055% | | 10.9 ± 0.6 | 23.8 ± 0.9 | 100.0 ± 1.3 |
| ***koLhcb*** | | **peak 669-672 nm** | | **peak 685-687 nm** | **peaks 732+750 nm** |
| | ctrl | 1.0 ± 0.8 | | 7.9 ± 1.0 | 100.0 ± 1.1 |
| | + α-DM 0.0055% | 1.2 ± 0.3 | | 20.1 ± 0.4 | 100.0 ± 1.2 |

**Supplementary Figure S4. Analysis of 77K fluorescence emission spectra of thylakoids from WT and *LHC* mutants.** (A) Gaussian deconvolution of the 77K emission spectra of thylakoids from wild type and mutants, before (*upper panels*) and after (*lower panels*) supplement of α-DM 0.0055%. Spectra were normalized to the max emission of PSI-LHCI at 733 nm. λexc = 440 nm. Max λ of the Gaussians used are reported. (B) Values (amplitudes) of the Gaussian decomposition of the fluorescence spectra reported in panel A.

### Supplementary Tables

**Supplementary Table S1.** Synopsis of LHC composition of the genotypes used in this work.

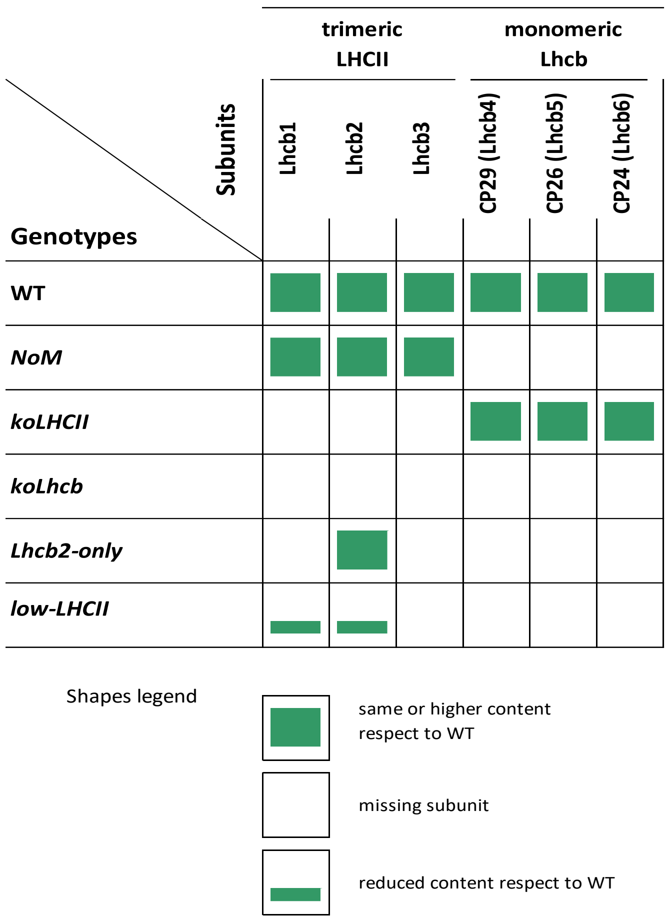

**Supplementary Table S2.** Sequences of sgRNA used for genome editing.

| Target | sgRNA |
|---|---|
| ***Lhcb1.1 - Lhcb1.5*** | GGTTCACAGATCTTCAGCGA |
| | ATGGACCC AAGTACTTGACT |
| | TGTGGATAACTTCT AGCTCA |
| | GGCTACTCAAGTTATCCTCA |
| | GAAGCGGCCGTGTGACAATG |
| | AGAAGTTATCCACAGCAGGT |
| | GAGGACTTGCTTTACCCCGG |
| | AGGGGAGGAGAGAGCCATTG |
| ***Lhcb2.1 - Lhcb2.3*** | CGCAAGGTTGGTGTATCCGG |
| | TGGAGGGCTTGAGAGCCGTT |
| | CGCCCAAGACGCCCACCTTT |
| | AGACTTGACGGTACGACGCA |
| | AGGGCTTGACCCGCTTTACC |
| | GGATCAAGTTAGGGTTTCCG |
| | CCAACATTGCCCATCTACTG |
| | TCTGGGCTGTTCAAGTTGTG |

**Supplementary Table S3. Lhcb1 and Lhcb2 content of *Arabidopsis* lines *lowLHCII* and *Lhcb2-only*.** The abundance of Lhcb subunits was determined on independent lines by SDS–PAGE fractionation of thylakoid proteins and immunotitration with specific primary antobodies. Lhcb content was normalized (norm.) to that of the CP47 (PSII core) subunit. Data are expressed as mean ± s.d., $n = 4$ biologically independent samples. Values marked with different letters are significantly different from each other within the column (ANOVA followed by Tukey's post-hoc test at a significance level of $P < 0.05$).

| | Protein content (norm. to WT level) | |
|---|---|---|
| | **Lhcb1** | **Lhcb2** |
| WT | 1.000 ± 0.046 [a] | 1.000 ± 0.152 [a] |
| *lowLHCII* L1 | 0.192 ± 0.033 [b] | 0.082 ± 0.026 [b] |
| *lowLHCII* L2 | 0.320 ± 0.096 [b] | 0.077 ± 0.016 [b] |
| *Lhcb2-only* | nd | 1.439 ± 0.218 [c] |

**Supplementary Table S4. Photosynthetic pigment composition of leaves from wild type and mutant lines.** Pigments were extracted from dark-adapted leaves with 85% acetone, separated and quantified by HPLC. Data are normalized to 100 Chl *a* + *b* molecules and are expressed as mean ± SD, n = 6 biologically independent samples.

| | Carotenoid content (normalized to 100 Chl *a* + *b*) | | | |
|---|---|---|---|---|
| | **Neoxanthin** | **Violaxanthin** | **Lutein** | **β-Carotene** |
| **WT** | 3.7 ± 0.2 | 3.0 ± 0.4 | 12.8 ± 0.3 | 7.7 ± 0.2 |
| ***NoM*** | 4.0 ± 0.1 | 3.7 ± 0.4 | 13.9 ± 0.2 | 7.5 ± 0.2 |
| ***koLHCII #17*** | 1.6 ± 0.2 | 5.1 ± 0.2 | 11.6 ± 0.3 | 11.4 ± 0.4 |
| ***koLHCII #36*** | 1.8 ± 0.3 | 5.3 ± 0.4 | 11.6 ± 0.5 | 11.3 ± 0.1 |
| ***koLhcb #9*** | 0.7 ± 0.2 | 6.4 ± 0.3 | 11.2 ± 0.2 | 13.8 ± 0.3 |
| ***koLhcb #16*** | 1.0 ± 0.2 | 6.1 ± 0.4 | 11.7 ± 0.1 | 13.7 ± 0.3 |
| ***ch1*** | 1.6 ± 0.6 | 6.3 ± 0.6 | 14.6 ± 1.0 | 13.6 ± 0.7 |
| ***ch1 koLhcb5*** | 2.0 ± 0.8 | 6.0 ± 1.3 | 14.8 ± 0.5 | 16.2 ± 1.3 |
| ***lowLHCII #14*** | 2.3 ± 0.1 | 4.8 ± 0.4 | 11.7 ± 1.5 | 10.2 ± 2.0 |
| ***lowLHCII #32*** | 1.8 ± 0.6 | 5.1 ± 0.8 | 11.5 ± 1.8 | 12.3 ± 1.1 |
| ***Lhcb2-only*** | 1.8 ± 0.3 | 4.8 ± 0.5 | 11.3 ± 2.2 | 10.7 ± 1.8 |

**Supplementary Table S5. PS stoichiometry and Chl fluorescence induction parameters determined for leaves of wild type and mutants.** PSI/PSII ratios were determined by immunotitration analysis with primary antibodies directed against the PSII core subunit PsbB (CP47) and the PSI core subunit PsaA. $F_m/F_0$ and $F_v/F_m$ values were obtained by analysis of chlorophyll fluorescence measured at the Dual-PAM. The connectivity parameter J was determined from fast Chl fluorescence induction in DCMU-treated leaves, measured with green light (7 µmol $m^{-2}$ $s^{-1}$). Values are shown as mean ± SD (biologically independent samples n = 3 for PSI / PSII ratio measurements, n ≥ 5 for Chl fluorescence analysis). Values for *Mantoniella squamata* were reported in (Trissl & Wilhelm, 1993).

| | PSI / PSII (PsaA / CP47) | $F_v$ / $F_m$ | $F_m$ / $F_0$ | J |
|---|---|---|---|---|
| **WT** | 1.00 ± 0.04[a] | 0.82 ± 0.01[a] | 5.56 ± 0.31[a] | 2.05 ± 0.24[a] |
| ***NoM*** | 1.00 ± 0.06[a] | 0.61 ± 0.01[b] | 2.56 ± 0.07[b] | 0.79 ± 0.13[b] |
| ***koLHCII #17*** | 0.46 ± 0.07[b] | 0.76 ± 0.01[c] | 4.17 ± 0.17[c] | 1.76 ± 0.07[a] |
| ***koLHCII #36*** | 0.45 ± 0.04[b] | 0.76 ± 0.01[c] | 4.17 ± 0.17[c] | 1.71 ± 0.10[a] |
| ***koLhcb #9*** | 0.66 ± 0.07[c] | 0.50± 0.02[e] | 2.00 ± 0.08[e] | 0.58 ± 0.25[b] |
| ***koLhcb #16*** | 0.70 ± 0.05[c] | 0.54 ± 0.03[e] | 2.17 ± 0.14[e] | 0.60 ± 0.39[b] |
| ***Mantoniella squamata (Trissl & Wilhelm, 1993)*** | 0.4 - 0.65 | 0.5 - 0.6 | 1.5 - 2.5 | nd |
